# Solubility, Hydration, and Sulphate Coordination in Cubic Insulin Crystals from Esrapid^TM^ Monomers Stabilized in Divalent Anionic Form of Citric Acid

**DOI:** 10.1101/2025.03.06.641839

**Authors:** Esra Ayan, Hiroaki Matsuura, Yoshiaki Kawano, Zain Abhari, Abdullah Kepceoğlu, Ahmet Katı, Takehiko Tosha, Hasan Demirci

## Abstract

Insulin’s structural adaptations have been extensively studied at neutral and basic pH; however, the effects of water and anion coordination on allosteric regions under various acidic conditions remain unexplored. Given the critical role of polar interactions in allosteric modulation, investigating structured water movements across extended pH intervals is essential for understanding solvent-mediated stabilization mechanisms in the monomer form of insulin. Structures of acid-stable cubic insulin crystals were determined in the divalent anionic form of citric acid solutions over a pH range of 2 to 6 to investigate the effects of water and anion coordination, along with charge distribution, on protein conformation. Synchrotron X-ray diffraction data were collected at resolutions ranging from 1.4 to 1.76 Å, with refined models exhibiting *R*-factors between 0.19 and 0.21. While the spatial arrangement of most proteins is highly conserved, ∼90% of the water coordination network and polar interactions alter local residue motion and intrinsic dynamics of the structures as the pH is changed. This is in line with previously determined structures at pH 7-11, allowing for a comprehensive structural analysis in the pH range of pH 2–11. Three key observations emerged: **(i)** water coordinations undergo a prominent shift toward the isoelectric point of insulin (pH 5-7), **(ii)** water molecules and anions function as allosteric modulators to stabilize the T-state of insulin at the pH range 2 to 6, and **(iii)** at extreme pH values 2 and 11, increased solubility correlates with the structural adoption of insulin’s most active form, wherein hydration within the allosteric pocket supports monomer stabilization in the T-state. Combined with the computational analyses, pH-dependent electrostatic redistributions primarily affect side-chain dynamics and local protein motion. This solvent-coupled allosteric regulation provides a mechanistic framework for solvent-mediated protein stabilization, offering a novel insight into the rational design of insulin formulations through controlled protonation and hydration strategies.

## Introduction

The first crystallographic structure of insulin was determined in 1969, revealing a hexameric organization where three identical dimers were arranged around two zinc ions [1]. Metal ions directly coordinate B10 histidine residues across the three dimers, contributing to the stabilization of the hexameric assembly. Within each monomer of the dimeric unit, the A chain adopts a conformation with two α-helices, while the B chain features one α-helix and two extended β-strand-like segments. This conformation corresponds to the T-state of insulin, which represents its highest activity in the monomer form of insulin [2]. The second hexameric form of insulin provided direct structural evidence for its intrinsic conformational flexibility. Under high chloride ion concentrations, the allosteric region of three B chains (residues B1–B8) undergoes partial unfolding and adopts a frayed α-helical conformation, currently termed the R^f^ state [3]. In the presence of phenol during crystallization, six allosteric sites adopt a fully extended α-helical conformation, characteristic of the R-state [4]. This structural transition does not disrupt hexamer assembly; instead, it enhances hexamer stability progressively by promoting tighter packing, stabilizing the R-state of the allosteric pocket, and ligand-induced cooperativity [2].

Under acidic conditions, insulin crystallizes in monomeric form instead of hexameric form, as protonation of the B10 histidine side chains prevents coordination with metal ions, and allosteric regions remain T-state to provide activity of insulin [5]. Similarly, insulin remains monomeric at pH values above 7 in the absence of zinc ions [6]. The cubic crystal structure of the monomer form of insulin, which belongs to the highest-symmetry space group, was previously determined using the molecular replacement method [7] and refined to 1.7 Å resolution with an *R* factor of 17.3% [8]. Structural studies on cubic insulin crystals have primarily focused on investigating the effects of varying physicochemical parameters, including pH, ionic strength, specific ionic species, and water activity, since monomer form has great interest in insulin activity mechanisms. Multiple cubic insulin crystal structures have been systematically analyzed under diverse conditions, such as in 1 M Na₂SO₄ solutions across a pH range of 5–11 [9], in carbonate-based solutions containing various monovalent cations at pH 9.5–10 [10], and in highly concentrated 2 M glucose solutions [11]. The cubic insulin structure shows high structural stability and remained largely conserved across these conditions. Even in cases where conformational changes induced notable shifts in atomic positions, the overall integrity of the insulin molecule and crystal lattice remained largely undisturbed.

Gursky et al. (1994) [9] investigated stereospecific dihaloalkane binding in cubic insulin crystals and observed a conformational shift in GluB13 toward lowering the pH from 9.0 to 5.0 in 1 M Na₂SO₄. Additionally, sulfate ions were coordinated by rearranged PheB1 NH₃⁺ groups. Gursky et al. (1992) also conducted a detailed analysis of cubic insulin crystals in 0.1 M sodium salt solutions across a pH range of 7–11 [6]. However, the effects of the range of low pH levels on cubic insulin crystals remain largely unexplored. Studies on insulin fibrillation have demonstrated that acidic conditions promote fibril formation of insulin, with the rate of fibrillation being dependent on the type of acid used (H₂SO₄ > HCl > H₃PO₄ > acetic acid) [12, 13]. To reduce the fibrillation propensity of monomeric insulin, we substituted the asparagine residue at the C-terminus of chain A with glycine due to its enhanced solubility under acidic conditions [14, 15]. Additionally, citric acid, a triprotic solubilizing agent with superior buffering capacity and mild pH-regulating properties [16, 17], was utilized to stabilize the insulin monomer and was further incorporated into the crystallization buffer to maintain structural integrity.

To investigate the structural and dynamic transitions of insulin at various acidic conditions, we have determined six crystal structures within the acidic pH range of 2.0 to 6.0. The crystallographic titration of cubic insulin crystals will elucidate probabilistic phenomena, such as transient coordination shifts in the water-anion dynamics surrounding the allosteric pocket in monomeric insulin. This study will provide deeper insights into the effects of pH on the dynamic behavior of polar residues in monomeric insulin and its effect on anion binding to biomacromolecules.

## Method

### Production of acid-stable insulin monomer

Esrapid^TM^ insulin monomer (hereafter referred to as mIns) was produced in our previous study [18]. Briefly, the recombinant mIns gene was cloned into the pET28a(+) vector and subsequently introduced into the E. coli BL21 Rosetta-2 strain for high-level expression. Independent bacterial cultures, each harboring the recombinant mIns gene, were cultivated in 6 liters of either LB-Miller medium or Terrific Broth (TB). Both media were supplemented with 35 μg/mL chloramphenicol and 50 μg/mL kanamycin to maintain selective pressure. The cultures were incubated at 37°C with continuous agitation at 110 rpm using a New Brunswick Innova 4430R shaker until they reached an OD_600_ of approximately 0.7. Protein expression was induced by adding Isopropyl β-D-1-thiogalactopyranoside (IPTG) to a final concentration of 0.4 mM, followed by further incubation at 37°C for 4 hours to promote recombinant protein production. Cells were then harvested at 4°C using a Beckman Allegra 15R centrifuge at 3500 rpm for 20 minutes. The successful expression of the recombinant protein was verified via precast TGX Mini-Protean gradient SDS-PAGE (Bio-Rad).

### The solubilization process of mIns

To achieve early purification and ensure the complete solubilization of inclusion bodies, 1 gram of mIns-containing cell pellets was individually resuspended in 10 mL of lysis buffer consisting of 25 mM Tris-HCl (pH 8.0) and 5 mM ethylenediaminetetraacetic acid (EDTA). The resuspended cells were then homogenized and subjected to sonication for cell lysis. The resulting lysate was then centrifuged at 6000 × g for 7 minutes at 4°C to remove cell debris. The obtained pellets were washed with a series of buffers to further refine inclusion bodies. Initially, the pellets were resuspended in wash buffer A, which contained 25 mM Tris-HCl (pH 8.0), 5 mM EDTA, 0.1% (v/v) Triton X-100, and 1 M urea. This was followed by resuspension in wash buffer B, composed of 25 mM Tris and 2 M urea. Each suspension underwent sonication for 30 seconds in an ice bath, followed by centrifugation at 8000 × g for 20 minutes at 4°C. To further isolate the inclusion bodies, the remaining cell debris was resuspended at a concentration of 0.1 g/mL in a binding buffer containing 25 mM Tris-HCl (pH 8.4), 5 mM 2-mercaptoethanol (BME), and 8 M urea. The suspension was then subjected to centrifugation at 10,000 × g for 30 minutes at 4°C, effectively recovering inclusion bodies enriched with fusion proteins. Finally, the resulting supernatants were filtered using a 0.2-micron Millipore filter to ensure purity and remove any residual particulates.

### Purification, refolding, and tryptic digestion process of mIns

The sulfitolysis of mIns was performed at 25°C for 4 hours using 200 mM sodium sulfite (Na_2_SO_4_) and 20 mM sodium tetrathionate (Na₂S₄O₆). Following this reaction, manual gravity-based Ni-NTA affinity chromatography was employed. The column was first equilibrated with one column volume of equilibration buffer, composed of 25 mM Tris-HCl (pH 8.0), 5 mM 2-mercaptoethanol (BME), and 8 M urea. Protein elution was carried out using a buffer containing 25 mM Tris-HCl (pH 8.0), 5 mM BME, 8 M urea, and 250 mM imidazole. At each purification step, protein purity was assessed through 20% SDS-PAGE gel electrophoresis, followed by Coomassie staining. High-purity fractions were collected and subsequently filtered using a 0.2-micron Millipore filter. The mIns refolding process was conducted in accordance with the previously established protocol [19]. The purified proinsulin (0.5 mg/mL) was subjected to dialysis against a buffer containing 25 mM Tris-HCl (pH 9.0), 8 M urea, and 0.5 mM EDTA at 4°C. To remove potential aggregates, the sample was centrifuged at 10,000 × g for 30 minutes at 4°C. The supernatant was then dialyzed into a refolding solution consisting of 0.1 M Glycine/NaOH (pH 10.5), 0.5 M urea, 0.5 mM EDTA, and 5% glycerol, with continuous stirring at 15°C overnight. Refolded mIns underwent a subsequent dialysis step in 25 mM Tris-HCl (pH 8.0) supplemented with 5% glycerol for 18 hours at 4°C. For proinsulin cleavage, tryptic digestion was performed using trypsin at a 1:1000 (v/v) ratio for 4 hours at 30°C. After proteolysis, pH-based precipitation was induced by adding 20 mM citric acid to the protein solution. The precipitated protein was then recovered by centrifugation at 3500 rpm for 5 minutes at 4°C, and the resulting pellet was resuspended in 20 mM citric acid to quench trypsin activity and facilitate its removal. The solubilized mIns was filtered through a 0.2 μm pore diameter filter and further purified via size-exclusion chromatography (SEC) using 20 mM citric acid buffer. High-purity fractions were collected, followed by a second pH precipitation step using 1 M Tris (pH 10.0). The INSv pellet was then solubilized in 20 mM citric acid buffer, and its concentration was adjusted to 9 mg/mL in the total volume of 10 mL.

### Proliferative effect of insulin monomers in hiPS cell culture

Human-induced pluripotent stem (hiPS) cell lines were generated from peripheral blood mononuclear cells (PBMCs) using an integration-free Sendai virus to introduce the Yamanaka factors [20]. The resulting cell lines were expanded on Matrigel in mTeSR1 medium (StemCell Technologies) and routinely passaged with 0.5 mM EDTA. Before experiments, all cells were thawed and acclimated to the TeSR-E8 medium for at least two weeks. The mIns analog, as described in the methods section, was successfully synthesized. For experiments involving insulin treatment, hiPS cells were cultured in a controlled incubator under an E8 medium supplemented with key growth-supporting factors. This medium contained 100 U/ml penicillin, 100 μg/ml streptomycin, 20 mg/ml transferrin, 3 mM sodium selenite, 100 mg/ml ascorbic acid, 1 mg/ml FGF2-G3, 0.5 μg/ml TGF-β3, and 20 mg/ml of either the novel mIns analog or commercially available insulin. Cells were maintained at 37°C under 5% CO₂. Cellular growth changes were systematically monitored at defined time points— day 1, day 3, day 5, day 7, and day 9—using the Mateo TL Digital Transmitted Light Microscope. The mIns insulin analog exhibited a growth-promoting effect on human iPS cells comparable to that of porcine insulin

### Crystallization of mIns insulin analog

To facilitate crystallization, monomer samples in 20 mM citric acid were subjected to screening using the oil-based sitting drop vapor diffusion micro-batch technique. A comprehensive initial crystallization screen was conducted by testing approximately 3000 commercially available sparse matrix and grid screen conditions [21]. For the screening process, equal volumes (0.83 µL, v/v, 1:1 ratio) of the crystallization solutions and 18 mg/mL mIns solution were mixed within a 72-well Terasaki plate (Cat#654,180, Greiner Bio-One, Austria). Each well was then sealed with 16.6 µL of paraffin oil (Cat#ZS.100510.5000, ZAG Kimya, Türkiye) and incubated at 4°C. The formation of crystals within the wells of the Terasaki plates was then monitored using a compound light microscope. Following the crystallization optimization, the protocol was adapted from a batch method to facilitate the formation of the cubic crystal form of insulin within a pH range of 2–6. To achieve precise pH control, 20 mM citric acid or 0.5 M sodium phosphate dibasic dihydrate was utilized, while 0.1 M sodium sulfate was employed as the precipitant. Crystals were refined using the vapor diffusion sitting drop technique. Each crystallization drop comprised a 1:1 (500 µL:500 µL) mixture of insulin solution (18 mg/mL mIns analog dissolved in 20 mM citric acid at pH 2) and reservoir solution (0.1 M sodium sulfate adjusted to pH 2 with citric acid), with pH levels continuously monitored using a micro pH meter. The resulting zinc-free cubic crystals of the mIns analog were further adjusted using 0.5 M sodium phosphate dibasic dihydrate, with pH systematically varied between 2 and 6 to explore its impact on crystallization dynamics. During pH 7 and so on, crystals gradually disappeared, probably due to the lattice order being disrupted.

### Sample delivery and data collection

X-ray diffraction data of a single crystal at diverse pH values was collected from the BL32XU beamline at SPring-8 (Japan) utilizing the automated data acquisition system *ZOO* [22]. Prior to data collection, the crystals were cryogenically preserved by rapid freezing in liquid nitrogen. Then, they were stored in pucks and transferred to a dewar filled with liquid nitrogen within the beamline hutch. Data acquisition was conducted using a continuous helical scanning method with a total oscillation range of 360°. The experimental parameters were as follows: an oscillation step size of 0.1°, an exposure time of 0.02 seconds per frame, and a beam dimension of 10 µm (horizontal) × 15 µm (vertical). The X-ray wavelength was set to 1 Å, while the average radiation dose per crystal volume was 10 MGy. Data collection was performed at a cryogenic temperature of 100 K using an EIGER X 9M detector (Dectris). The acquired datasets were subsequently processed automatically using XDS within the KAMO pipeline.

### Structure determination and refinement

The monoclinic crystals of mIns was resolved within the I123 space group using the automated molecular replacement algorithm *PHASER* [23], incorporated within the *PHENIX* software suite [24]. The structure determination process was initiated with a previously published X-ray model (PDB ID: 5VIZ), which served as the starting search model. The atomic coordinates from 5VIZ were first subjected to rigid-body refinement within *PHENIX* as an initial step. The following refinement procedures involved simulated annealing, positional optimization of atomic coordinates, and Translation/Libration/Screw (TLS) refinement to enhance model accuracy. Additionally, a composite omit map approach, available in *PHENIX*, was employed to pinpoint potential side chain modifications and water molecule positions. Model inspection and adjustments were carried out in *COOT* [25], with particular emphasis on regions exhibiting significant differences in electron density. Any spurious water molecules outside well-defined electron density regions were manually removed. The refined X-ray crystal structure was visualized and analyzed using *PyMOL* [26] and *COOT*.

### GNM analysis, calculation of hinges, and cross-correlations

The Gaussian Network Model (GNM) normal mode analysis was conducted using the *ProDy* package [27] to investigate the dynamical properties of six newly determined cryogenic-temperature mIns structures at various pHs. The analysis was performed utilizing only Cα atoms to generate contact maps. A Kirchhoff matrix (N × N, where N represents the total number of residues) was constructed and subsequently decomposed into normal modes. A cutoff distance of 8.0 Å was applied to define long-range interactions, yielding N − 1 nonzero normal modes. To compare global and local motions between the structures, square fluctuations were computed across the weighted slowest modes, which collectively accounted for 30% of the total variance, as well as for the ten fastest modes. Additionally, cross-correlations between residue fluctuations and hinge site analysis were derived from GNM modes to further elucidate their dynamic behavior.

### Principal Component Analysis

#### Data Collection and Preprocessing

The atomic coordinates of the six newly determined cryogenic-temperature mIns structures at various pHs (2, 3, 4, 5, 6, 7, 9, 10, 11) were extracted from the refined X-ray crystal structures. Each structure was parsed to extract the coordinates of the alpha carbon (Cα) atoms. *Alignment and Truncation.* To ensure consistency across all structures, the number of Cα atoms was standardized. The minimum number of Cα atoms present in any structure was determined, and all structures were truncated to this length by retaining only the first N Cα atoms, where N is the minimum number of Cα atoms across all structures. This step ensured that each structure had the same number of atoms for subsequent analysis. *Principal Component Analysis.* PCA was performed on the aligned and truncated Cα coordinates to identify the major modes of structural variation. The coordinates of each structure were flattened into a single vector, resulting in a matrix (X) of size (M×3N), where (M) is the number of structures and (3N) represents the x, y, and z coordinates of the N Cα atoms. *The PCA was conducted using the following steps:* **(i)** Mean Centering: The mean of each coordinate dimension (x, y, z) was subtracted from the corresponding coordinates to center the data. **(ii)** Covariance Matrix Calculation: The covariance matrix (C) of the centered data was computed. **(iii)** Eigen Decomposition: The eigenvalues and eigenvectors of the covariance matrix were calculated. The eigenvectors represent the principal components (PCs), and the eigenvalues indicate the variance explained by each PC. **(iv)** Projection: The centered data were projected onto the principal components to obtain the principal component scores. *Variance Explained.* The variance explained by each principal component was calculated as the ratio of the corresponding eigenvalue to the sum of all eigenvalues. This ratio was expressed as a percentage to indicate the proportion of total variance captured by each principal component. *Visualization.* The first two principal components were plotted to visualize the major modes of variation among the insulin structures. Each structure was represented as a point in the PC1-PC2 plane, with labels indicating the pH value. Additionally, a bar plot of the variance explained by each principal component was generated to illustrate the contribution of each component to the overall variance. *Specific Atom Analysis.* To further investigate the structural variations, specific Cα atoms of interest were identified based on their residue number and chain ID. The coordinates of these atoms were extracted and analyzed to understand their contributions to the principal components. *Software and Tools.* All analyses were performed using MATLAB (MathWorks, Inc.). The *pdbread* function was used to parse PDB files, and the *pca* function was employed to perform the principal component analysis.

## Result and Discussion

### Quality of structures

We determined six high-resolution cubic crystal structures (pH 2 to 6) of a monomer insulin analog (mIns, esrapid^TM^) within the I2₁3 space group. The pH of the solution containing cubic mIns crystals was adjusted using the 20 mM citric acid or 0.5 M sodium phosphate dibasic dihydrate solution, where stabilization was obtained through the titration of protein functional groups and allosteric interactions mediated by sulfate coordination. Data collection statistics are summarized in **Table 1**. Accordingly, the asymmetric unit of the crystals consists of a single monomer composed of two cross-linked peptide chains (A and B). The electron density maps enabled the modeling of 44 residues, except for side chains of E4_A_, E21_B_, F25_B_, and K29_B_, which lacked detectable electron density (**Fig 1A-E**). Zinc-free insulin crystals contain a single SO₄²⁻ anion per monomer at the allosteric site (1-8 residues in chain B) and exhibit a progressive decrease in water molecules toward the isoelectric point (pH: 5-6) of mIns. To evaluate structural alterations among C-alphas as a function of ionic strength, we also incorporated PDB structures determined at pH 7 to 11. Superposition of all individual monomers yielded Root Mean Square Deviation (RMSD) values ranging from 0.170 Å to 0.306 Å, possibly in response to increasing positive-to-neutral ion charge in the pH range pH 2 to 6 (**Fig 1F-J**) and from 0.273 Å to 0.230 Å with neutral-to-negative ion charge in the pH range 7 to 11 (**Fig 1K-O**) under strong ionic strength conditions. According to the extended Debye–Hückel equation [28], ionic strength gradually increases from pH 2 to 6, with values of 1.33 M, 1.40 M, 1.63 M, 2.0 M, and 2.38 M, respectively, whereas it also progressively increases from pH 7 to 11, reaching 0.75 M, 0.875 M, 1.00 M, and 1.125 M, respectively. However, this shift in ionic strength induces only minor conformational variations among individual chains, suggesting minimal structural adaptability of monomers (**Fig 1P**). This is in line with the previously comparative studies of electron density difference maps and the refined atomic models at pH 5 to 9 [29] as well as pH 7 to 11 [6], which reveal that the spatial arrangements of the majority of protein residues and well-ordered solvent molecules remain largely consistent across these structures, exhibiting minimal positional deviations. As anticipated for high-resolution structures, the coordinate geometry of mIns structures is well-defined (Δr.m.s. chirality ranging from a minimum of 0.063 Å^3^ to a maximum of 0.072 Å^3^), all main-chain dihedral angles are confined within the allowed regions of the Ramachandran plot, ensuring structural integrity (**see, Table 1**).

**Figure 1.**
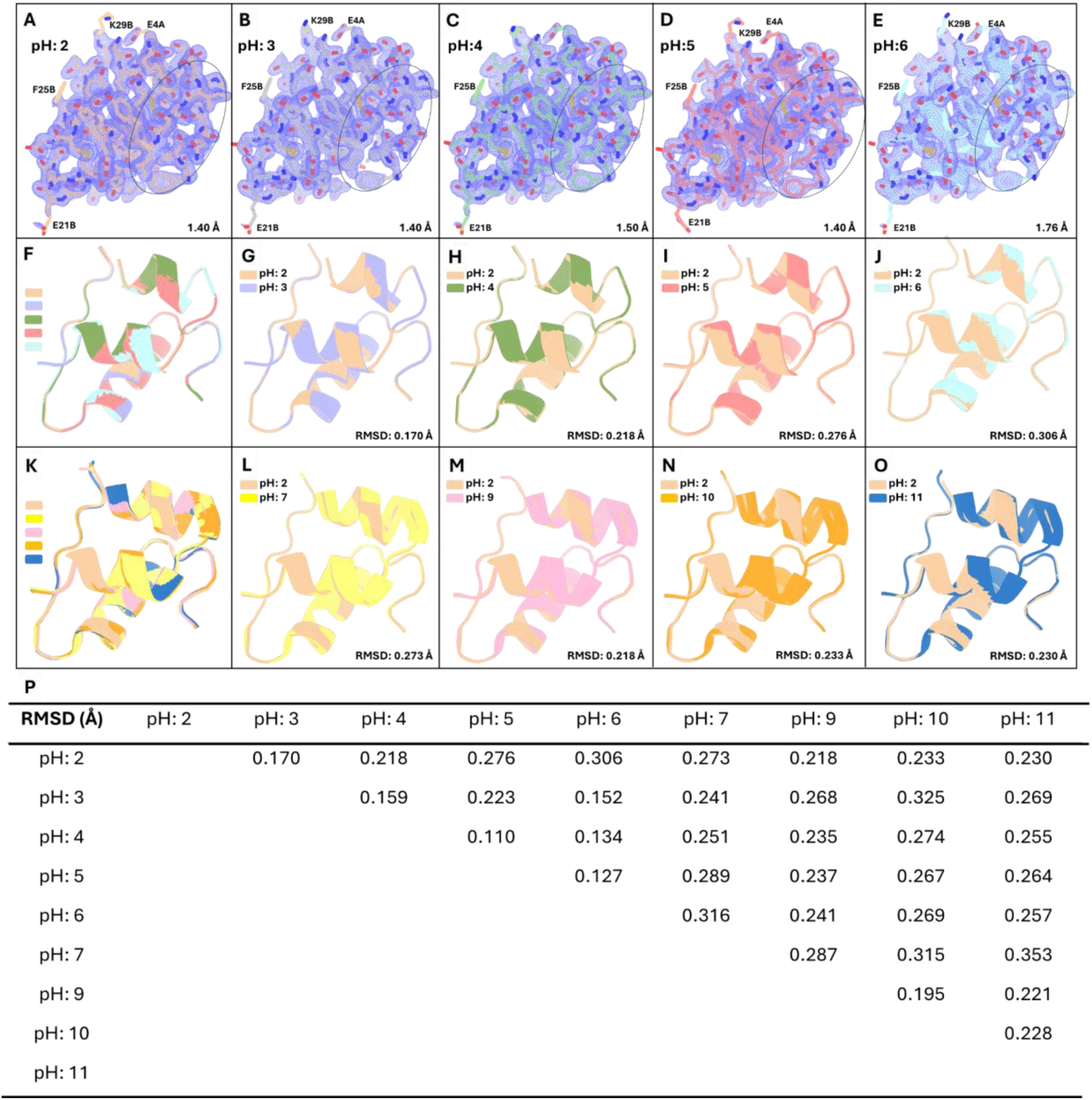
Superposition of monomer insulins in the pH range 2-11. **(A-E)** The structures of mIns are 2*Fo-Fc* simulated annealing-omit map at 1 sigma level, which is colored in slate. Residues E4A, E21B, F25B, and K29B are labeled as they exhibit a lack of electron density in those side-chain conformations. The allosteric site has been indicated as an elliptic dash line. **(F)** Superposition of each mIns structure in the pH range 2 to 6, showing minor conformational changes. **(G-J)** Superposition between pH 2 vs pH 3, 4, 5, 6, rising 0.170 Å to 0.306 Å RMSD values. **(K)** Superposition of each monomer structure in the pH range 7 to 11, showing minor conformational changes. **(L-O)** Superposition between pH 2 vs pH 7, 9, 10, 11, with RMSD values ranging from a minimum of 0.218 Å to a maximum of 0.273 Å. **(P)** Superposition between pH 2, 3, 4, 5, 6 vs pH 7, 9, 10, 11, with RMSD values ranging from a minimum of 0.110 Å to a maximum of 0.353 Å.

**Table 1.** Data collection and refinement statistics.

| Data collection | mIns (pH:2) | mIns (pH:3) | mIns (pH:4) | mIns (pH:5) | mIns (pH:6) |
| --- | --- | --- | --- | --- | --- |
| Beamline | Spring-8<br>(BL32XU) | Spring-8<br>(BL32XU) | Spring-8<br>(BL32XU) | Spring-8<br>(BL32XU) | Spring-8<br>(BL32XU) |
| Resolution (Å) | 18.38 - 1.4<br>(1.451 - 1.4) | 18.39 - 1.4<br>(1.45 - 1.4) | 18.39 - 1.5<br>(1.554 - 1.5) | 19.55-1.4<br>(1.45-1.4) | 19.57-1.76<br>(1.823 - 1.76) |
| Space group | I 21 3 | I 21 3 | I 21 3 | I 21 3 | I 21 3 |
| Unit cell | 77.97 77.97<br>77.97 90 90 90 | 78.02 78.02<br>78.02 90 90 90 | 78.01 78.01<br>78.01 90 90 90 | 78.18 78.18<br>78.18 90 90 90 | 78.26 78.26<br>78.26 90 90 90 |
| No of reflections | 197102 | 195778 | 196717 | 198771 | 198628 |
| Unique reflections | 15684 (1581) | 15644 (1536) | 12802 (1253) | 15820 (1548) | 8078 (818) |
| Multiplicity | 48.0 (25.0) | 48.0 (25.0) | 48.0 (25.0) | 48.0 (25.0) | 48.0 (25.0) |
| Completeness % | 99.89 (99.75) | 99.49 (97.71) | 99.88 (99.76) | 99.90 (99.81) | 99.86 (99.76) |
| Mean I/sigma(I) | 23.03 (1.14) | 35.36 (1.05) | 29.05 (0.78) | 16.97 (0.53) | 11.74 (0.28) |
| Wilson B-factor | 25.08 | 27.63 | 28.54 | 28.31 | 33.85 |
| CC1/2 | 1 (0.313) | 0.999 (0.724) | 1 (0.565) | 0.999 (0.508) | 0.998 (0.201) |
| CC* | 1 (0.69) | 1 (0.916) | 1 (0.85) | 1 (0.821) | 1 (0.579) |
| R-meas | 0.02072<br>(0.232) | 0.01996<br>(1.108) | 0.01444<br>(1.306) | 0.02459<br>(1.695) | 0.059<br>(10.48) |
| R-work | 0.1907<br>(0.3946) | 0.1964<br>(0.3933) | 0.1966<br>(0.3424) | 0.2142<br>(0.3818) | 0.2153<br>(0.3345) |
| R-free | 0.2270<br>(0.4125) | 0.2114<br>(0.4492) | 0.2201<br>(0.4131) | 0.2150<br>(0.4046) | 0.2376<br>(0.3662) |
| No. of reflections in ref. | 15684<br>(1581) | 15644<br>(1536) | 12802<br>(1253) | 15819<br>(1548) | 8078<br>(818) |
| Protein residues | 50 | 50 | 50 | 50 | 50 |
| RMS (bonds) | 0.008 | 0.012 | 0.014 | 0.005 | 0.012 |
| RMS (angles) | 0.93 | 1.09 | 1.19 | 0.69 | 1.04 |
| Ramachandran favored (%) | 100.00 | 100.00 | 100.00 | 100.00 | 97.83 |
| Ramachandran allowed (%) | 0.00 | 0.00 | 0.00 | 0.00 | 2.17 |
| Ramachandran outliers (%) | 0.00 | 0.00 | 0.00 | 0.00 | 0.00 |
| Average B-factor | 35.24 | 34.72 | 35.20 | 36.37 | 41.54 |
| macromolecules | 33.41 | 33.91 | 34.64 | 36.08 | 41.44 |
| ligands | 1 | 1 | 1 | 1 | 1 |
| solvent | 49.60 | 45.46 | 43.99 | 43.14 | 42.70 |

The average temperature factors for all main-chain and side-chain atoms of mIns are well-defined, ranging from a minimum of 34.72 Å² at pH 3 to a maximum of 41.54 Å² at pH 6, consistent with the overall Wilson plot values, which range from 25.08 Å² at pH 2 to 33.85 Å² at pH 6 (**Fig S1**). The lowest Wilson B-factor (25.08 Å²) and average B-factor (35.24 Å²) at pH 2 indicates the highest degree of structural rigidity (**Fig S1I-J**), while the progressive increase in both values toward pH 6 (**Fig S1A-H**), might indicate local atomic displacement, primarily attributed to translational disorder due to slight expansion in unit cell parameters [30]. This is also mutually supportive in validating ellipsoid representations of atomic displacement parameters and residue-specific fluctuations in the scatter plots, offering a quantitative validation of the observed local atomic displacement (**Fig S2**). The panels **K–S in Figure S1**, representing previously determined structures at pH 7–11, lack precise *B*-factor data for direct comparison; however, based on the trend observed in panels **A–J**, it is likely that flexibility continues to increase toward pH 7 before stabilizing or decreasing at extreme alkaline conditions. This observation aligns with a previous study investigating the pH-dependent effects on cubic insulin crystals within the pH range of 5–9 [29]. Notably, the average *B*-factor exhibits a biphasic trend, reaching its lowest values at both acidic and basic conditions (47.47 Å² at pH 5 and 48.34 Å² at pH 9) while peaking at 58.42 Å² at pH 6.98. This pattern reflects a progressive increase in atomic displacement parameters as the system approaches the isoelectric point, suggesting heightened structural flexibility near pH neutrality

The following section will be divided into two parts: **(i)** Analyzing the coordination of water molecules and allosteric binding of sulfate doubly charged anions in cubic insulin crystals as a function of pH and ionic strength while exploring the relationship between alterations in the polar network and the titration of protein functional groups. **(ii)** Investigating the causal interrelations between water molecules and mIns molecules through the Gaussian Network Model (GNM) and Principal Component Analysis (PCA), with a particular emphasis on the allosteric site dynamics, to elucidate the role of intrinsic residual mobility of cubic insulin crystals. Namely, both static binding interactions and dynamic effects are being investigated.

### Water coordination and sulfate ion binding

The overall polar interactions within mIns structures progressively decrease from pH 2 to pH 6 **(Fig 2A-J)**, whereas they increase from pH 7 to pH 11 **(Fig 2K-S)**. Specifically, the number of polar contacts at pH 2, pH 3, pH 4, pH 5, and pH 6 are 451, 435, 428, 418, and 407, respectively, while at pH 7, pH 9, pH 10, and pH 11, the values increase to 494, 480, 498, and 503, respectively. Notably, mIns at pH 2 **(Fig 2I-J)** and pH 11 **(Fig 2R-S)** exhibit the highest number of coordinated water molecules, which interact with allosteric residues and promote the stabilization of the T-state in mIns through a cooperative network of dipole-dipole and dipole-charge interactions. Conversely, those allosteric residues of monomer insulin correlate with a progressive reduction in the number of coordinated water molecules and polar contacts as the system approaches the isoelectric point (pH 5–7) **(Fig 2A-D and Fig 2K-L)**. This observation aligns with the established solubility profile of insulin, which is highest at both extreme pH values (pH 2 and pH 11) [31, 32]. These findings suggest that coordinated water molecules in the allosteric site, along with the polar interaction network, play a critical role in solvent-mediated cooperativity. Over pH 2 to 6, the sulfate ion engages in a network of charge-assisted hydrogen bonds with two critical allosteric residues of Phe1_B_ and Asn3_B_, supporting local conformational (T-state) stability **(Fig 2B, D, F, H, J)**. Sulfate ions are incorporated into both the protein structure and the crystal lattice, where they neutralize positive charges, contributing to protein stability and promoting crystallization. Their interactions cause local deformations, suggesting increased structural plasticity at low pH. Sulfate ions exhibit selective binding, preferring structurally stable regions over mobile positive charges; thus, they play a key role in salt bridge formation [5].

**Figure 2.**
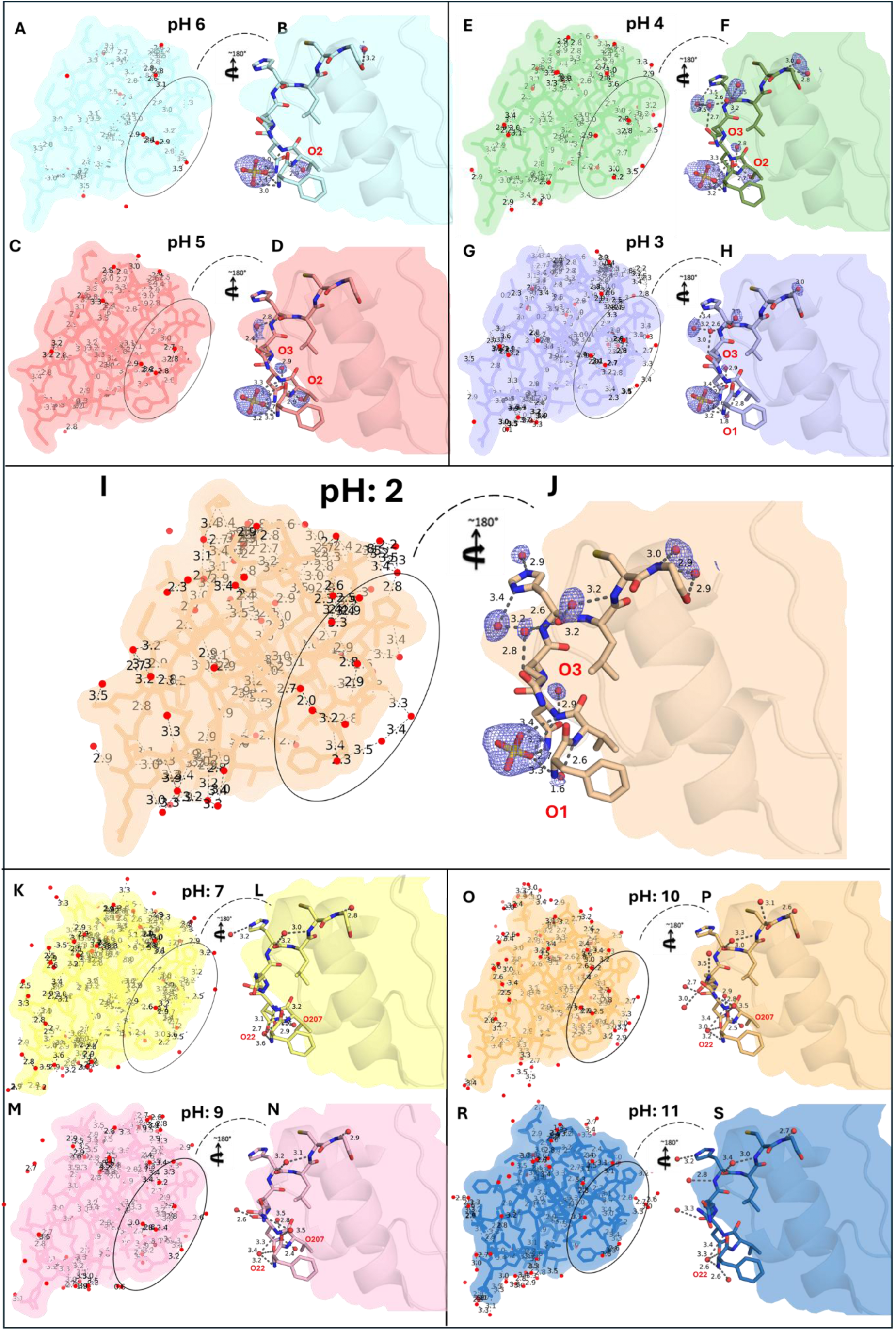
Overall polar interactions and coordination of structured water molecules within the allosteric pocket were reduced toward the isoelectric point of insulin. The overall and allosteric site residues of T-state monomer insulin structures were observed with **(A)** 407 polar interactions and **(B)** 2 water molecules, stabilized by SO₄ coordination at pH 6. At pH 5, the T-state monomer insulin structures exhibited **(C)** 418 polar interactions and **(D)** 3 water molecules, stabilized by SO₄ coordination. At pH 4, the T-state monomer displayed **(E)** 428 polar interactions and **(F)** 8 water molecules, stabilized by SO₄ coordination. Similarly, at pH 3, the overall and allosteric site residues of the T-state monomer exhibited **(G)** 435 polar interactions and **(H)** 6 water molecules, stabilized by SO₄ coordination. At pH 2, the overall and allosteric site residues involved **(I)** 451 polar contacts and **(J)** 9 water molecules, stabilized by SO₄ coordination. Likewise, the overall polar interactions were **(K)** 494, **(M)** 480, **(O)** 498, and **(R)** 503 in the pH range of 7–11 (PDB IDs: 1APH, 1BPH, 1CPH, and 1DPH), respectively, accompanied by **(L)** 5, **(N)** 7, **(P)** 10, and **(S)** 7 water coordinations toward higher pH values (pH 7 to 11). The stabilization of the T-state in porcine insulin monomers was provided by critical water coordinations (O22 and O207) instead of SO₄ stabilization at the pH range 7-11. Collectively, overall polar interactions and water coordinations were reduced toward the isoelectric point of the insulin monomer at various pH intervals, suggesting that an apparent solubility and hydration effect was observed at pH 2–3 and pH 10–11. All polar interactions were observed up to 3.6 Å. The allosteric site of the overall monomer indicated an elliptic dashed line.

Despite the literature [29], clear electron density indicates the presence of the sulfate ion even toward higher pHs, and refined occupancy of the sulfate ion remains constant at 1.00, indicating full site occupancy and no significant alterations in binding stoichiometry **(Fig S3).** The tetrahedral coordination geometry of sulfate ion serves as an H-bond acceptor through its electronegative oxygen atoms. In the pH range 2 to 6, the O atom of the sulfate closest to the threefold axis is modulated by symmetry-related N-terminal α-amino group of Phe1 through two H-bonds (∼3.0 Å and ∼3.2 Å), forming a distorted tetrahedral arrangement with local planarity in the binding network **(Fig 2B, D, F, H, J)**. The phenyl ring of Phe1 may engage in weak van der Waals interactions and potentially weak anion-π interactions with sulfate [33, 34], but the dominant stabilization arises from sulfate-mediated H-bonding with backbone amides. In this vein, previous studies examining the pH 7–11 range [6], where sulfate was not introduced, adopted a distinct allosteric stabilization mechanism. Water molecules coordinated with Phe1_B_ and Asn3_B_, effectively modulating T-state stabilization through a series of H-bond networks in the absence of anion coordination **(Fig 2L, N, P, S)**. This highlights that divalent anions might act as potential allosteric stabilizers for active monomer modulation (**Fig. S4**), as their bivalency enables participation in multivalent H-bonding [35] - similar to water coordination - thereby contributing to T-state stabilization effectively.

Likewise, dipole-charge electrostatic interactions (∼2.7 Å) with the side-chain amide (-CONH_2_) of symmetry-related Asn3 also support the allosteric regulatory network of insulin. Besides, Asn3 is involved in secondary H-bonds (∼2.9 Å) with critical water molecules, further stabilizing the local electrostatic environment **(Fig 2B, D, F, H, J)**. Asn3 residue is highly crucial, playing a dual role as an allosteric regulator and a key electrostatic contributor, forming coordinated interactions along the threefold symmetry axis of hexamer insulin [18, 36, 37]. Further stabilization of residues Phe1 to Asn3 is facilitated by certain water molecules (O1, O2, and O3). At pH 2, the interatomic distances between the nearest water molecule (O1) and the sulfate sulfur (S) are 3.3 Å **(Fig 2J; Fig S5A)**, which slightly contracts to 3.4 Å at pH 3 **(Fig 2H; Fig S5B)**. As the pH surpasses 4.0, O1 disappears within the coordination sphere, suggesting pH-dependent solvent reorganization. Thus, the S atom transitions to a more distant hydration shell, where its nearest water ligand (O2) is positioned at 7.8 Å at pH 4 and pH 6 **(Fig 2B, F; Fig S5C, E)**, slightly shifting to 7.9 Å at pH 5 **(Fig 2D; Fig S5D)**. This progressive displacement of structured water molecules suggests a pH-driven adaptation in hydration dynamics [38] and sulfate coordination geometry.

As the pH modulates the protonation states of key residues, the electrostatic landscape gradually shifts, affecting both sulfate coordination and hydration shell stability (**Fig 3**). Toward lower pHs, protonation-driven charge neutralization attenuates electrostatic contrast (**Fig 3A-J**), potentially constructing water-mediated stabilization. Conversely, toward higher pHs, deprotonation also attenuates electrostatic contrast (**Fig 3K-S**), providing neutralization and modulating solvent accessibility. This pH-dependent redistribution of electrostatic forces not only governs hydration dynamics but also modulates the extent of cavity formation (**Fig S6**). Calculation of the cavity area and volume displays a pH-dependent trend, peaking at pH 6 (Area: 20.3 Å², Volume: 6.8 Å³) and decreasing at both lower and higher pH values. In the acidic range (pH 2 to pH 6), cavity size gradually increases (pH 2: 4.8 Å² & 0.6 Å³ to pH 5: 7.2 Å² & 1.1 Å³ to pH 6: 20.3 Å² & 6.8 Å³), reflecting enhanced electrostatic polarization (**Fig 3A, B**) and decreases in water binding (**Fig 2A, B**) [39]. Conversely, in the alkaline range (pH 7 to pH 11), cavities shrink progressively (pH 7: 11.7 Å² & 2.6 Å³ to pH 9: 9.4 Å² & 2.1 Å³ to pH 11: 3.5 Å² & 0.3 Å³), suggesting that extensive deprotonation allows structured water reorganization (**Fig 2K-S**), promoting solvent-mediated cavity stabilization [40, 41].

**Figure 3.**
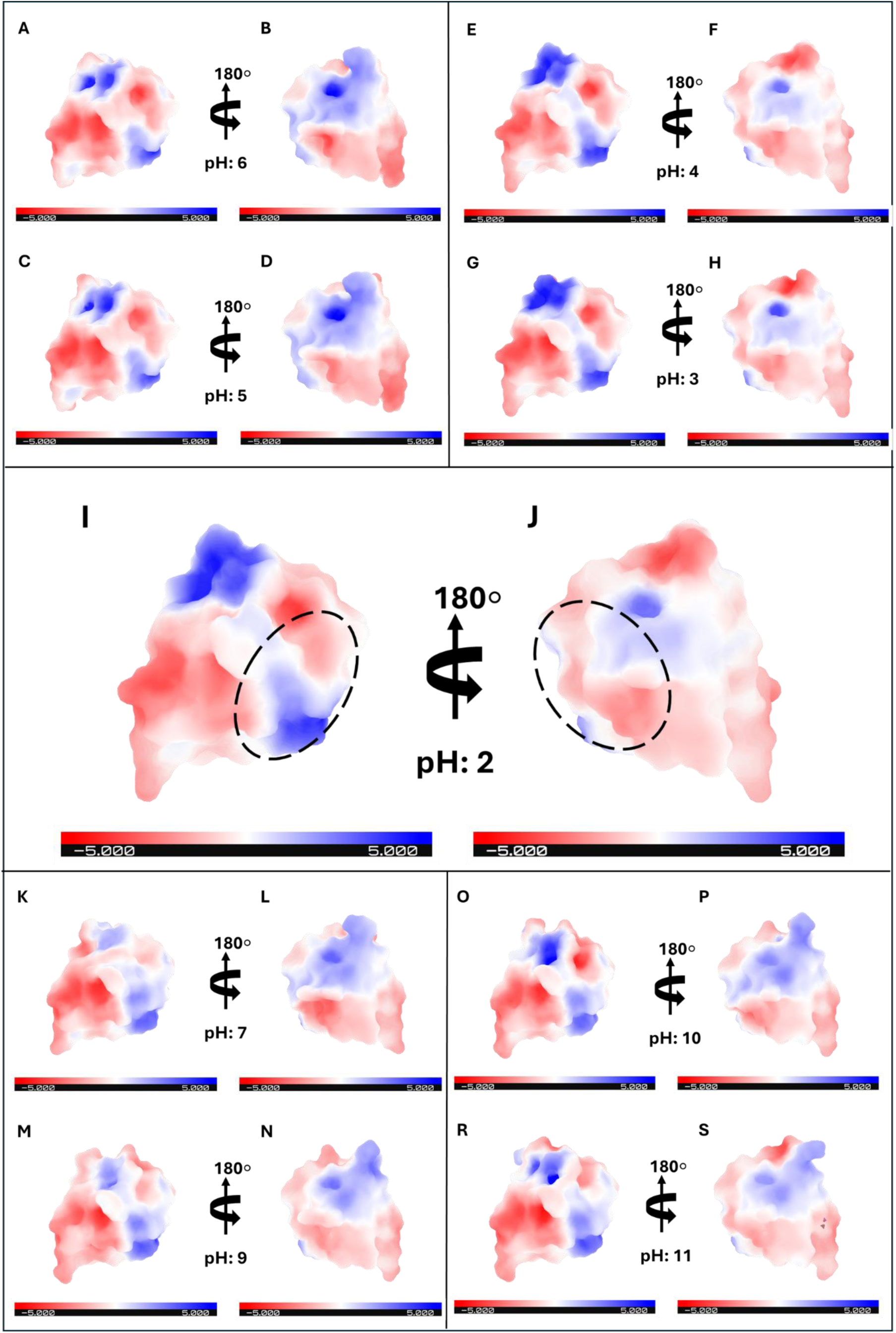
Comparison of electrostatic charge distribution in monomer insulin structures in the pH range 2-11. The color gradient represents electrostatic potential, with negative (basic) regions in blue and positive (acidic) regions in red. Each structure is shown in two orientations, rotated by 180°, to illustrate the charge distribution across the molecular surface. **(A-D)** The charge distribution displays a well-defined polarized pattern, with distinct negatively (blue) and positively (red) charged regions. The allosteric region maintains strong charge separation, indicative of a more native-like electrostatic profile. **(E-J)** The apparent shift in charge distribution is observed from pH 4 on, where the negative electrostatic potential is largely neutralized due to extensive protonation. The allosteric region (dashed ellipses) at pH 2, which previously contained strong negative patches, now appears predominantly neutral or weakly positive -due to the electrostatic surface potential of protonated basic groups can become more positive-, indicating a loss of electrostatic contrast and homogenization of surface charges. Emphasizes that a progressive neutralization of the insulin monomer’s surface charge as pH decreases, with the most substantial changes occurring between pH 5 and pH 2, particularly in the allosteric region, where negative charge patches are gradually lost due to protonation effects. Likewise, the progressive alteration in charge distribution becomes evident from pH 9 on **(K-N)**, as the positive electrostatic potential undergoes gradual neutralization due to extensive deprotonation **(O-S)**. At pH 11, the allosteric region, which previously exhibited prominent positive charge clusters, now appears largely neutral, indicating a reduction in electrostatic contrast and a more uniform surface charge distribution. This underscores a continuous decline in the asymmetry of the insulin monomer’s surface charge as pH increases, with the most prominent transformations occurring between pH 7 and pH 11, particularly in the allosteric region, where positive charge densities are systematically reduced through progressive deprotonation.

This bell-shaped distribution highlights the redistribution of electrostatic interactions and hydration dynamics in cavity formation, with pH 6 representing the peak cavity expansion state (**Fig S6**) due to enhanced electrostatic polarization (**Fig 3A, B**) as well as local structural heterogeneity (**Fig S1A, B; Table 1, Wilson *B*-factors**). Collectively, the coordination of sulfate binding, structured water molecules, and electrostatic interactions governs the allosteric stabilization of monomer insulin. Sulfate coordination with Phe1_B_ and Asn3_B_ establishes a rigid electrostatic framework, while the surrounding hydration shell modulates the local energy landscape, facilitating conformational stabilization.

### In silico structural dynamics

We examined the intrinsic dynamics of mIns structures (pH 2-6) and reference structures from PDB (pH 7-11) using the Gaussian Network Model (GNM), a coarse-grained elastic network model that determines the normal mode fluctuations of a given structure under equilibrium conditions [42, 43]. GNM is widely applied in protein dynamics studies, allowing for the characterization of both global dynamics, which are captured in the lowest frequency collective motions, and local high-frequency fluctuations, which define site-specific mobility profiles. Regions revealing significant displacement in low-frequency modes correspond to flexible segments, whereas those constrained in these modes are often part of hinge sites, which play critical roles in facilitating large-scale conformational changes [44, 45] or modulating allosteric communication [46]. Conversely, residues that undergo high-frequency fluctuations are typically referred to as *kinetically hot spots*, as they often contribute to folding nuclei or evolutionarily conserved interaction sites [47, 48].

Comparison of the fastest modes in the pH range 2-11 reveals that solvent coordination minimally affects the overall fluctuation profile but introduces distinct local modulations at specific residue positions, likely due to reorganizations of the coordinated water molecules (**Fig 4A-E**). In the acidic range (pH 2–6), the fluctuation profile between hydrated and non-hydrated structures remains largely consistent, except for pH 3 and pH 4 (**Fig 4B, C**), where an additional fluctuation peak emerges between residues 25–28 (Asn3_B_-Cys7_B_) in the presence of water molecules. This suggests that solvent interactions at those allosteric sites transiently alter the kinetic energy distribution, potentially due to water-induced dipole realignments or redistribution of H-bond networks [49]. Conversely, in the alkaline range (pH 7–11), a similar solvent-mediated modulation is observed at pH 11, where residues 38–42 (Leu17_B_-Glue21_B_) exhibit elevated mobility in the hydrated structure compared to other pH conditions (**Fig S7A-D**). This might indicate a shift in solvent accessibility and dipole realignment upon extensive deprotonation. The elevated kinetic activity of this segment suggests that hydration at pH 11 reorganizes the local stabilization networks [50, 51], probably influencing receptor binding capacity but not altering the fundamental positioning of kinetic hot spots. As we consider global motion over the pH range 2-11 with and without water coordination, we can discern minimal deviations in residue-specific displacements separated by hinges (**Fig 4F-J; Fig S7E-H**). Notably, hydrated structures exhibit suppressed flexibility, whereas non-hydrated structures display apparent fluctuations, suggesting a dampening effect of solvent coordination on large-scale dynamics. Despite the overall trend remaining consistent in the pH range 2-11, non-hydrated structures at pH 4–6 (**Fig 4H-J**) reveal a distinct pattern, identified based on local maxima in the mean-square fluctuation profiles, which predominantly align with the C-terminal regions of each chain. This suggests that the reduced water coordination toward pH 6 may influence hinge stabilization, potentially modulating inter-residue couplings [29, 52]. Notably, cross-correlation analysis of structured waters within the allosteric region reveals a significant interdependency between solvent coordination and allosteric dynamics in the pH 2–6 range (**Fig 4K-O**). The structured waters exhibit a strong positive correlation (close to red, cc > 0.5) with residues in the allosteric core. This correlation diminishes progressively as the pH increases (**Fig 4N-O**), in line with the observed decrease in solvent occupancy and allosteric coupling strength. The theoretical fluctuations calculated from all modes, with and without water bindings at pH range 2-11, were aligned to the experimental *B*-factors (**Fig 4P-Z; Fig S7I-P)**, which is compatible with the trend observed in ten slow modes (**Fig 4F-J**; **Fig 75E-H)**. The theoretical fluctuations showed a high correlation with the experimental *B*-factor at the selected cut-off distance of 8.0 Å: overall correlation coefficient ranging from 0.795 at pH 2 to 0.730 at pH 6 (**Fig 4P-Z**) and 0.635 at pH 7 to 0.698 at pH 11 (**Fig S7I-P**). Accordingly, the presence of the water molecules further stabilized the thermal vibrations in the hydrated structures (**Fig 4P-U; Fig S7I-L**) compared to non-hydrated ones (**Fig 4V-Z; Fig S7M-P**) at the pH range of 2-11. Overall findings suggest that structured water molecules act as integral modulators of allosteric transitions by mediating long-range dynamic couplings between functionally significant residues. This observation aligns with studies showing that oligomeric proteins achieve cooperative allostery via water-mediated polar networks that effectively mimic substrate interactions [53].

**Figure 4.**
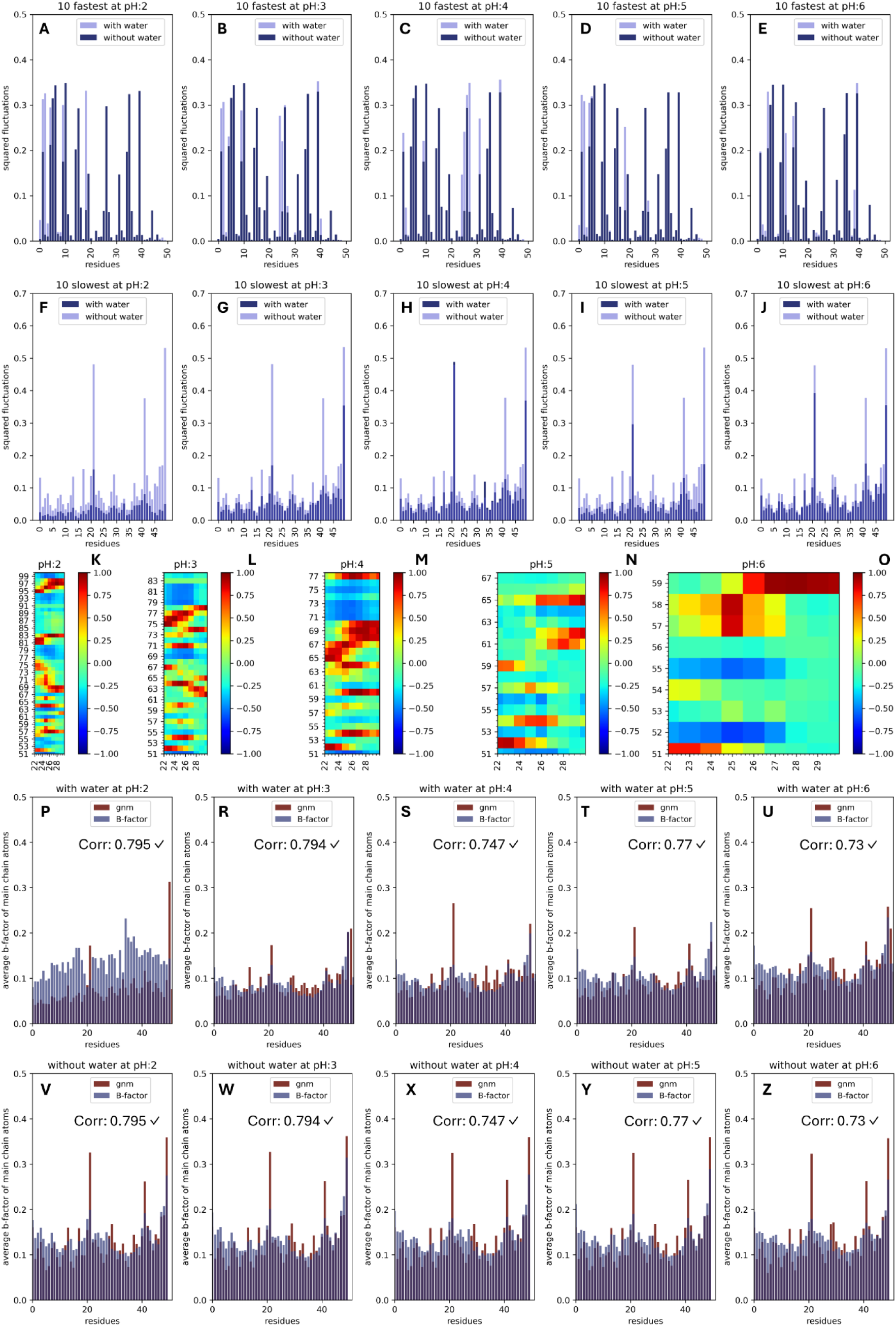
GNM analysis of mIns at the pH range 2-6. **(A–E)** Local protein motions from the ten fastest GNM modes, comparing structures with and without water coordination in the pH range of 2–6. The mean-square fluctuations of individual modes are plotted for each mIns residue. **(F–J)** Global protein motions from the ten slowest GNM modes. The weighted squared fluctuations of mIns, both with and without water coordination, are presented after normalization. **(K–O)** Cross-correlation analysis between allosteric residues (B1–B8) and all water molecules. The correlation matrix, calculated from the ten slowest GNM modes, maps residue motion correlations, where the x-axis represents allosteric residues and the y-axis specifies the entire water molecules. The color scale spans from −1.00 (strongly anticorrelated motions, blue) to 0.0 (uncorrelated, green) and 1.00 (highly correlated, red). **(P–U)** and **(V–Z)** shows the correlation between theoretical GNM fluctuations and experimental *B*-factors for each mIns variant, with and without water coordination, respectively.

Principal Component Analysis (PCA) was also performed to quantify pH-dependent variations at the pH range 2-11, identify key residues contributing to dynamic fluctuations, and determine the relationship between allosteric regions and the global motion profile in insulin monomers. PCA is a dimensionality reduction technique widely applied in protein dynamics to identify dominant motion patterns and structurally significant fluctuations [54]. By decomposing covariance matrices derived from atomic displacements, PCA reveals the most functionally relevant residue movements. Each principal component (PC) captures a progressively smaller variance, with the first principal component (PC1) accounting for the largest variance [55, 56]. The PCA scatter plot is depicted in **Figure 5A**. The separation of pH clusters along PC1 suggests that pH-dependent shifts are not random but follow a well-defined trajectory. Specifically, structures at pH 2–5 cluster closely together, with pH 6 slightly separated but still in proximity, forming one group. In contrast, structures at pH 7–10 form another distinct group, with pH 11 showing similar variance to the first group and being distinctly separated from the second group along PC2. This reflects a transition in the intrinsic dynamic profile of the monomer form, probably due to protonation-dependent electrostatic rearrangements and solvent reorganization. To unearth the key residues contributing to dynamic variability, PC1 residue contribution analysis was also performed (**Fig. S8A**), revealing that specific amino acids exhibit disproportionate control on the principal motion profile. Notably, residues from Chain A (Cys7, Thr8, Ser9, Ile10) correspond to sequence variations between porcine and human insulin, while residues from Chain B (Asn3, Gln4, His5, Leu6, Cys7) are localized within the allosteric region, suggesting a potential link between allosteric interactions and pH-mediated dynamic modulations. Despite the observed variance, no major conformational changes are detected (**Fig 5B-M; Fig S9**), indicating that the primary source of variation arises from local fluctuations rather than global structural rearrangements, which is in line with the ten fastest GNM modes with additional fluctuation peak emerging between residues 25–28 (Asn3_B_-Cys7_B_) in the presence of water molecules (**Fig 4A-E; Fig S7A-D**). This confirms that protonation-dependent electrostatic shifts and solvent interactions subtly modulate residue-specific collective motion without inducing distinct conformational movements.

**Figure 5.**
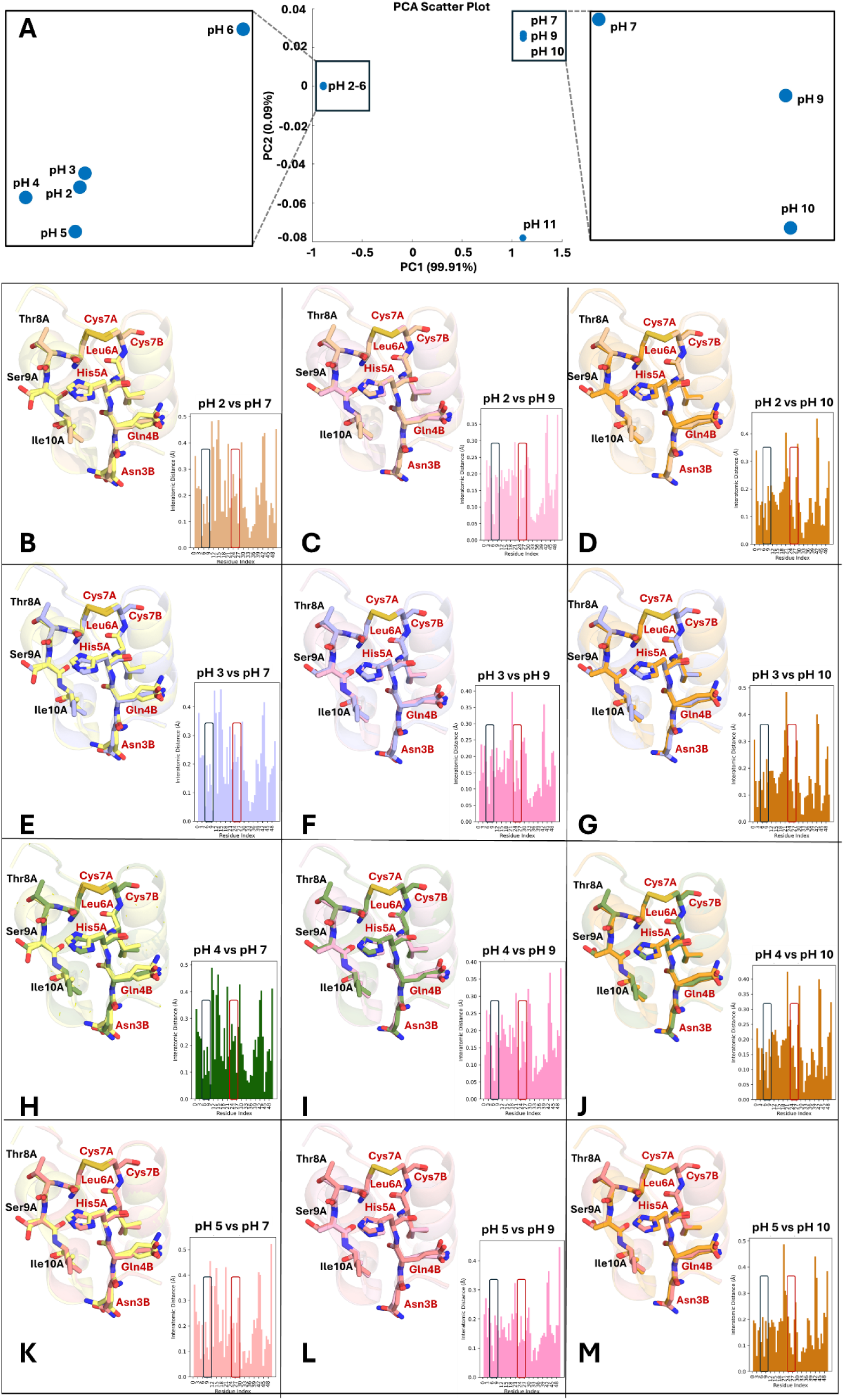
PCA analysis between pH 2-11 and interatomic distance between clusters calculated by PCA. **(A)** PCA across the pH range of 2 to 11 reveals two distinct clusters. The first cluster comprises samples at pH 2–5, which exhibit proximity to pH 6, while the second cluster includes samples at pH 7–10, closer to pH 11. These two subgroups demonstrate apparent variance from each other, indicating distinct physicochemical properties across the pH spectrum. PC1, accounting for 99.91% of the variance, represents the primary axis of differentiation, where pH 11 appears distinctly separated from the rest. PC2, explaining only 0.09% of the variance, captures minimal secondary variations. **(B-M)** Interatomic pairwise distance comparing structures from pH 2–5 (cluster 1) and pH 7–10 (cluster 2) reveals no major conformational alterations, as confirmed by structural alignment. Variant residues between porcine and human insulin are highlighted with a black rectangle, while allosteric sites are indicated with a red rectangle in the plots Key residues calculated by contribution analysis from PC1 (*see Fig S8*) are highlighted in stick representation onto structures: residues labeled in black indicate sequence variations between porcine and human insulin’s chain A, while those labeled in red emphasize allosteric residues revealing variations between cluster 1 and cluster 2. While no large-scale C-alphas reorganization is observed, the PCA-derived variance suggests that the differentiation between clusters is probably driven by alterations in side-chain dynamics, local protein motion, as well as changes in H-bonding networks and water coordination.

## Conclusion

This study provides **(i)** a comprehensive structural and computational analysis of pH-dependent (pH 2 to 6) allosteric water coordination and polar network reorganizations in cubic insulin crystals, revealing fundamental insights into the T-state stabilization mechanisms of the newly designed monomer form of insulin stabilized by divalent anion citric acid. Our findings demonstrate that hydration dynamics within the allosteric pocket exhibit a well-defined pH-dependent transition, which we validated through high-resolution electron density mapping, GNM, and PCA (**Fig 2**; **Fig 4**; **Fig 5**). This was also compared with reference insulin monomers in the pH range of 7–11, **(ii)** providing a comprehensive analysis of water coordination and the polar network across pH 2–11, where solubility and hydration increase toward extreme pH values (pH 2-3 and pH 10-11) but decrease near the isoelectric point (pH 5-7) of monomer insulin. This suggests that water coordination in the allosteric pocket at extreme pH values is closely related to the stabilization of the monomer in the T-state (active) form (**Fig S4**). Our results establish that **(iii)** anion coordination acts as a water-alternative stabilizing factor for the T-state monomer, indicating that anions serve as allosteric stabilizers, neutralizing all positive charges on the protein. This suggests a dual regulatory mechanism where water and anions cooperatively influence the intrinsic dynamics and thermodynamic stability of the T-status of insulin under varying pH conditions. **(iv)** PCA and GNM analyses further revealed that pH-dependent electrostatic redistributions primarily affect side-chain dynamics, local protein motion, and H-bonding networks rather than inducing backbone conformational transitions. This solvent-coupled allosteric modulation, which involves long-range electrostatic interactions and local conformational dynamics, offers a novel perspective on solvent-mediated protein stability. Furthermore, **(v)** the pH-dependent biphasic behavior of atomic displacement parameters suggests that monomeric insulin undergoes a hydration-driven shift in allosteric regulation, where structured water molecules act as integral modulators of cooperative dynamic networks in the absence of anion coordination even. The hydration shell reorganization observed in sulfate-free insulin crystals further supports the idea that solvent interactions alone can stabilize the T-state monomer, providing an alternative mechanism to anion coordination. This offers that in the absence of phenolic agents stabilizing the monomer in the R-state, water and anions act as opposing allosteric stabilizers, favoring the T-state form of monomer insulin. Beyond the fundamental report, these findings provide a structural framework for rational insulin formulation design, particularly in engineering stable and bioavailable insulin variants through controlled protonation and hydration strategies. The identification of anion-driven and hydration-driven allosteric stabilization mechanisms could pave the way for alternative formulation strategies that enhance insulin solubility, extend shelf life, and optimize pharmacokinetics, independent of traditional phenolic stabilizers.

## Data availability

The coordinates and structure factors have been deposited in the Protein Data Bank under the accession codes 9M4X (pH 2), 9M4Y (pH 3), 9M4Z (pH 4), 9M50 (pH 5), and 9M51 (pH 6). The raw dataset (.h5) can be accessed via the link: https://doi.org/10.5281/zenodo.14934063

## Acknowledgment

This work was supported by the SACLA Research Support Program for Graduate Students (proposals 2023B8058 and 2023B2761). The authors thank Kay Diederichs for numerous discussions on crystallographic methods and advanced refinement of Esrapid^TM^ structures, as well as for critically reviewing the manuscript. They also acknowledge Arda Odabaş for optimizing cell culture conditions for insulin treatment.

## Author Contributions

Esrapid^TM^ was designed, produced, purified, and characterized by E.A. Crystals were obtained and pH-optimized by E.A. Synchrotron data at SPring-8 was collected by E.A., Y.K., and T.T. Crystal data processing was performed by E.A. and H.M. Structures were refined and interpreted by E.A. GNM analysis was performed and interpreted by E.A. PCA analysis was calculated by Ab.K. The manuscript is written and reviewed by E.A., H.M., Y.K., Z.A., A.K., Ab.K., T.T., and H.D. All of the authors read and acknowledged the manuscript.

---

**Supplementary Figure 1.**
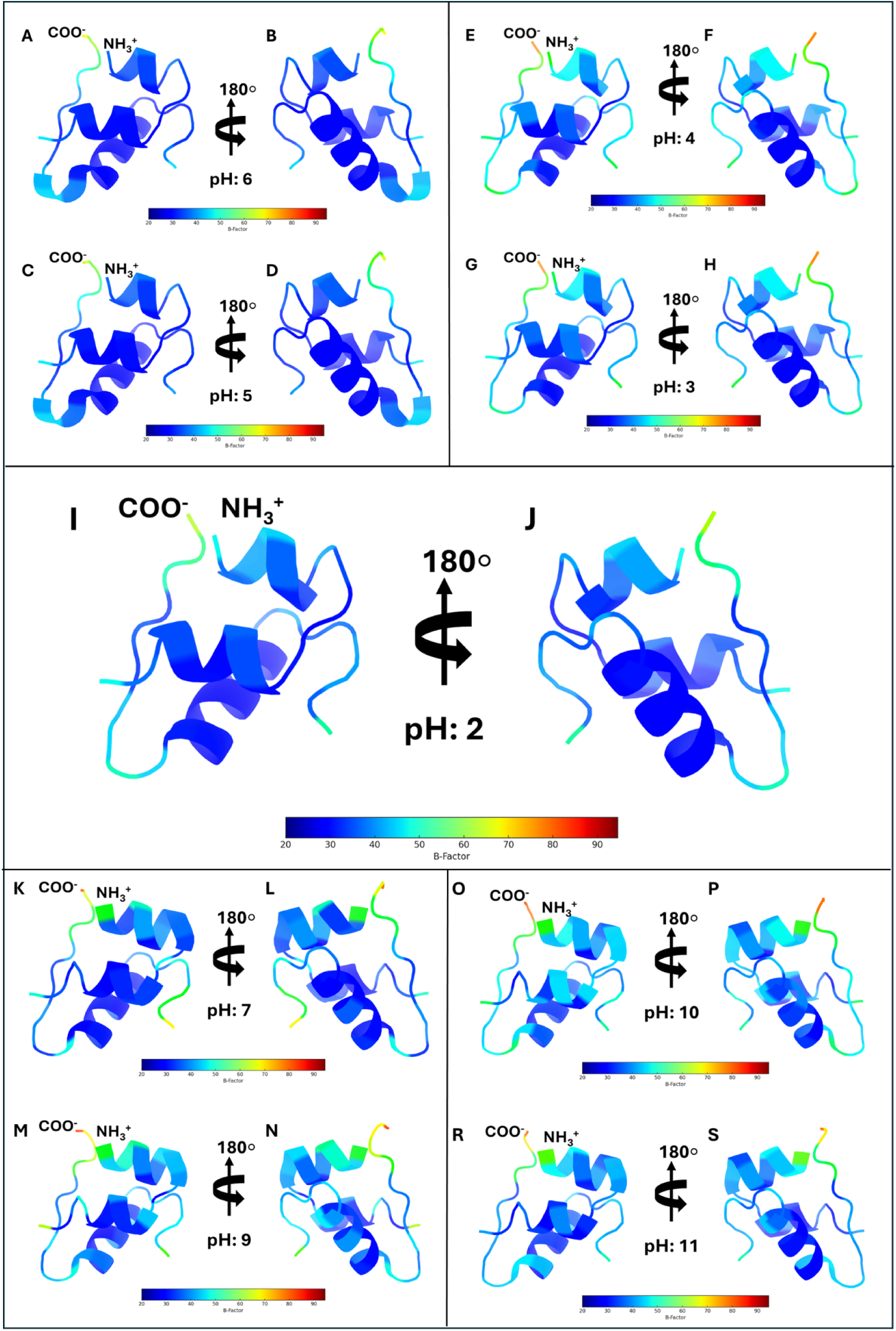
Cartoon representations of the *B*-factor distribution in monomer insulin structures across varying pH conditions (pH 2–11). The structures are color-coded according to the atomic displacement parameters, with the *B*-factor scale ranging from 20 Å² (blue, low flexibility) to 90 Å² (red, high flexibility). Each panel presents the insulin monomer in two orientations rotated by 180° to illustrate differential flexibility across the structure. **(A–B)** The highest *B*-factors among the low-pH structures are observed, particularly in loop regions and termini, indicating increased atomic displacement and local conformational heterogeneity as the system approaches the isoelectric point at pH 6. **(C–D)** A moderate increase in *B*-factors at pH 5 is evident compared to lower pH values, reflecting heightened thermal vibrations. **(E–F)** The distribution of atomic displacement at pH 4 remains relatively uniform, but loop regions begin to exhibit minor increases due to thermal vibrations or static disorder. **(G–H)** The *B*-factor values decrease compared to pH 4–6, indicating a more stabilized monomeric conformation. **(I–J)** The lowest average *B*-factors are observed at pH 2, suggesting maximal structural rigidity and reduced local atomic displacement. **(K–L)** *B*-factors at pH 7 begin to stabilize relative to pH 6, suggesting an adaptation to neutral pH conditions. **(M–N)** Structural rigidity increases, as evidenced by the shift of blue regions across the C-alphas backbone. **(O–P)** The structure exhibits further stabilization at pH 10, with decreasing *B*-factor fluctuations. **(R–S)** The *B*-factor at pH 11 distribution remains relatively low, indicating that high-pH conditions stabilize the monomeric structure similarly to low-pH conditions (pH 2–3).

**Supplementary Figure 2.**
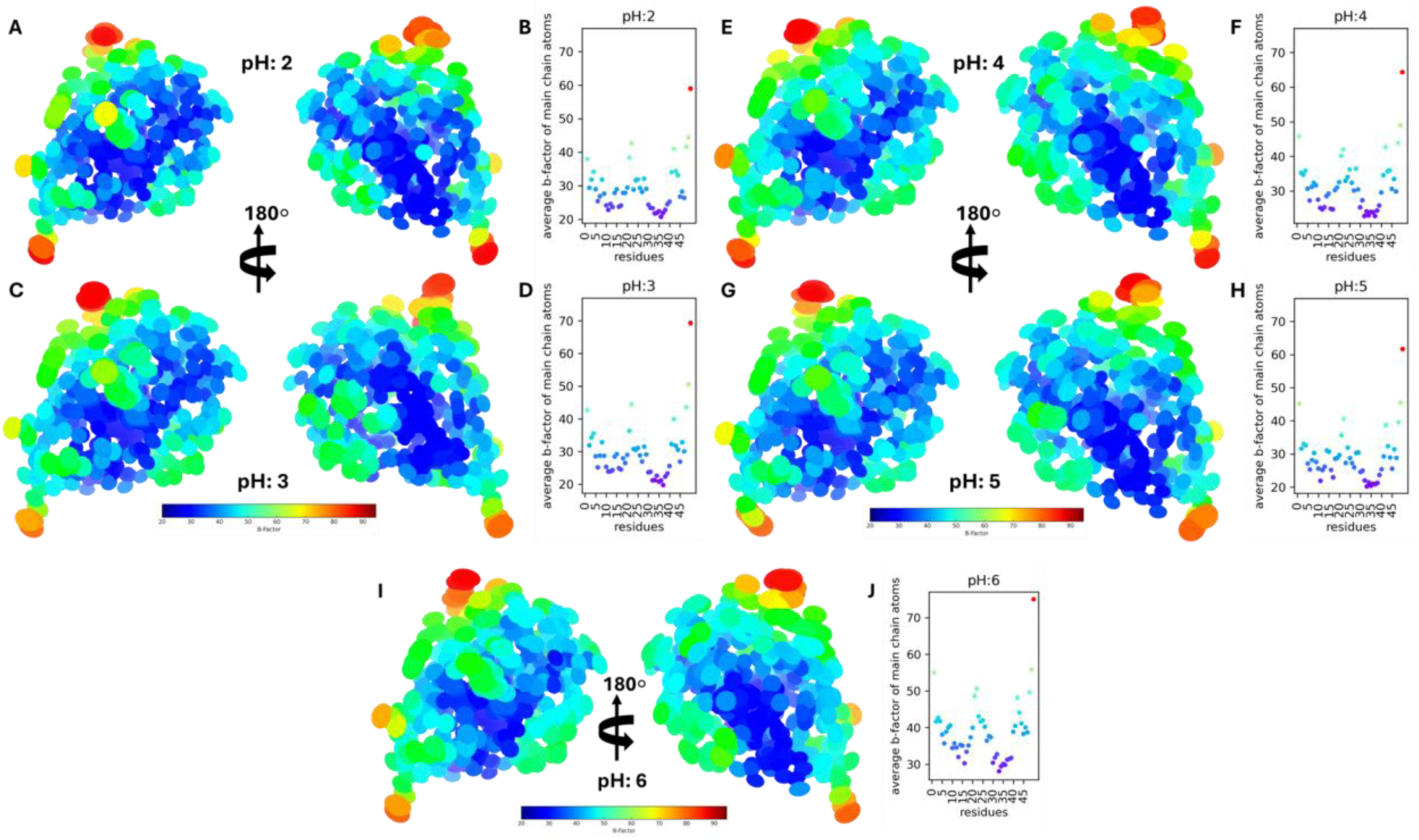
Detailed representation of average *B*-factors of monomeric insulin in the pH range 2–6. The average *B*-factor per residue is calculated by mapping onto the structure, with corresponding values displayed in scatter plots. Each panel **(A, C, E, G, I)** presents the anisotropic displacement parameters visualized as thermal ellipsoids, illustrating directionally dependent atomic displacements. The color-coded *B*-factor scale ranges from 20 Å² (blue, low flexibility) to 90 Å² (red, high flexibility). Panels **(B, D, F, H, J)** depict residue-specific average *B*-factors, reinforcing the trends observed in Supplementary Figure 1 (cartoon representation). Both representations exhibit a biphasic trend, with lower *B*-factors at pH 2 and 3, a gradual increase towards pH 6, and peak atomic displacement near the isoelectric point. The high *B*-factor regions localized at termini and loop regions remain consistent between the two figures, confirming the robustness of the structural flexibility distribution. This suggests that pH-dependent variations in atomic displacement are probably influenced by hydration effects and lattice-packing interactions, which is in line with electrostatic charge distributions of mIns in the pH range 2-6 (Fig 3).

**Supplementary Figure 3.**
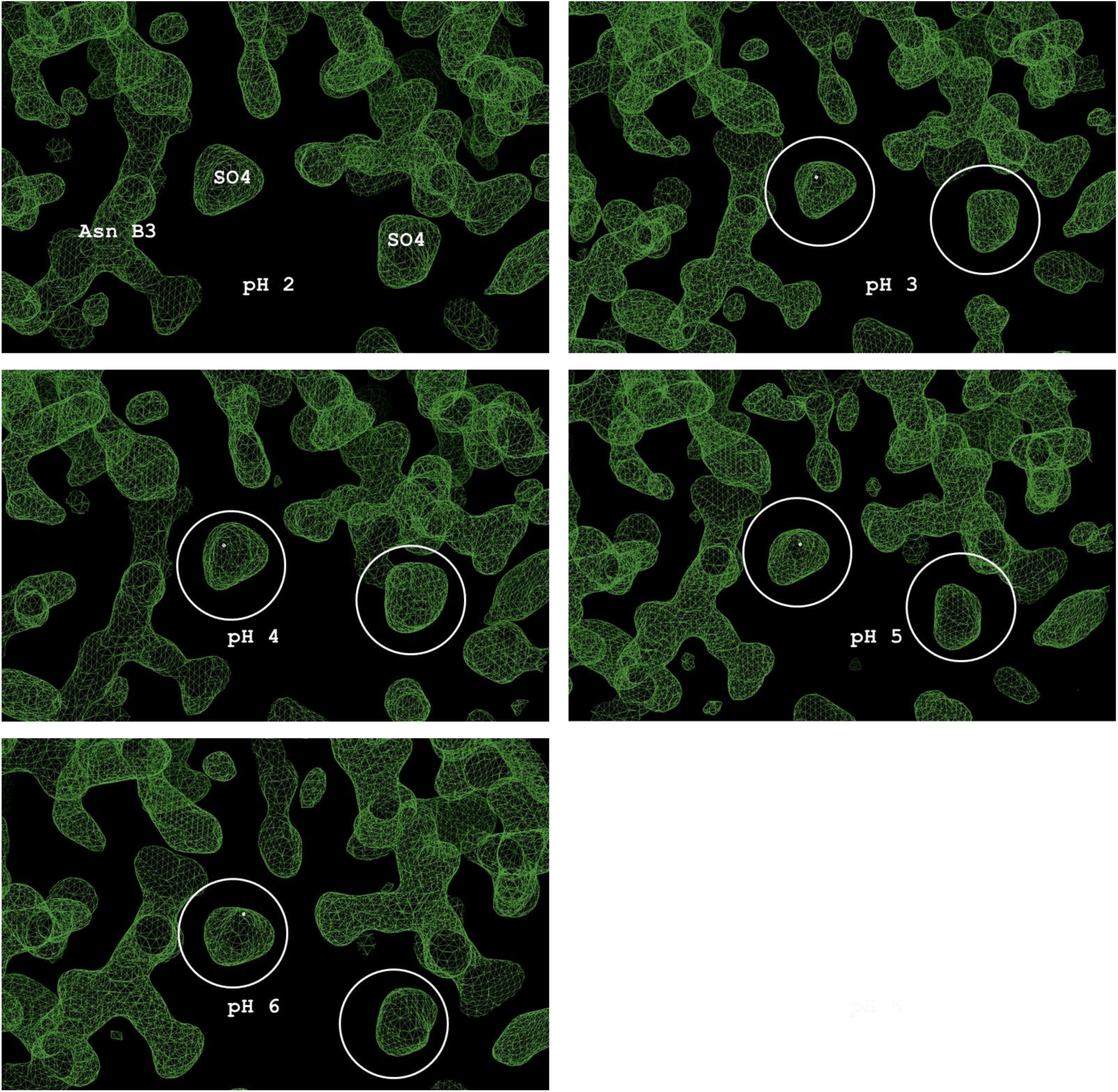
Structural models at five distinct pH values with corresponding 2*Fo-Fc* electron density maps. The structural region encompassing Asn3_B_ and the sulfate ion is displayed, with 2*Fo-Fc* maps contoured at 1σ to reflect full occupancy. Subfigures represent structures determined at pH 2, 3, 4, 5, and 6, respectively.

**Supplementary Figure 4.**
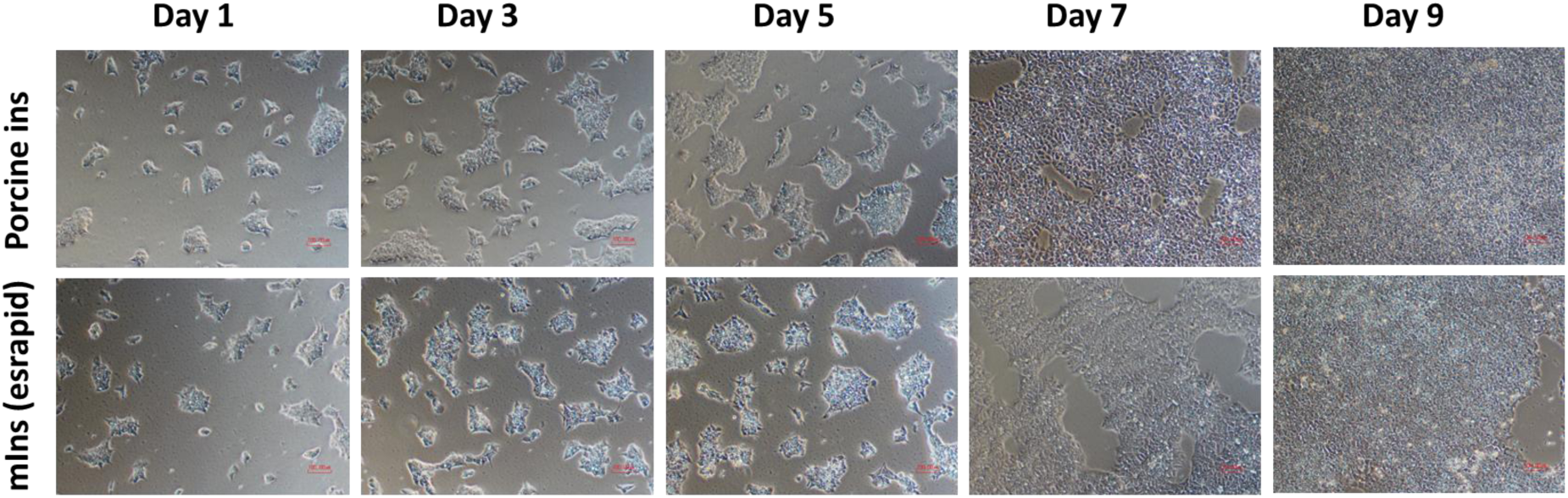
Proliferative effect of mIns and commercially porcine insulin molecule in cell culture. The t-state of the insulin monomer is the highest active form of insulin structure. As confirmed by structural analyses, T-state forms of mIns and porcine insulin molecules show their activity in the cell culture, promoting their proliferation. hiPS cells, which cannot grow in the absence of soluble insulin, were supplemented with 20 mg/ml acid-stable mIns analog stabilized in citric acid and 20 mg/ml commercially available porcine insulin during the time course. Comparatively, both promoted cell growth.

**Supplementary Figure 5.**
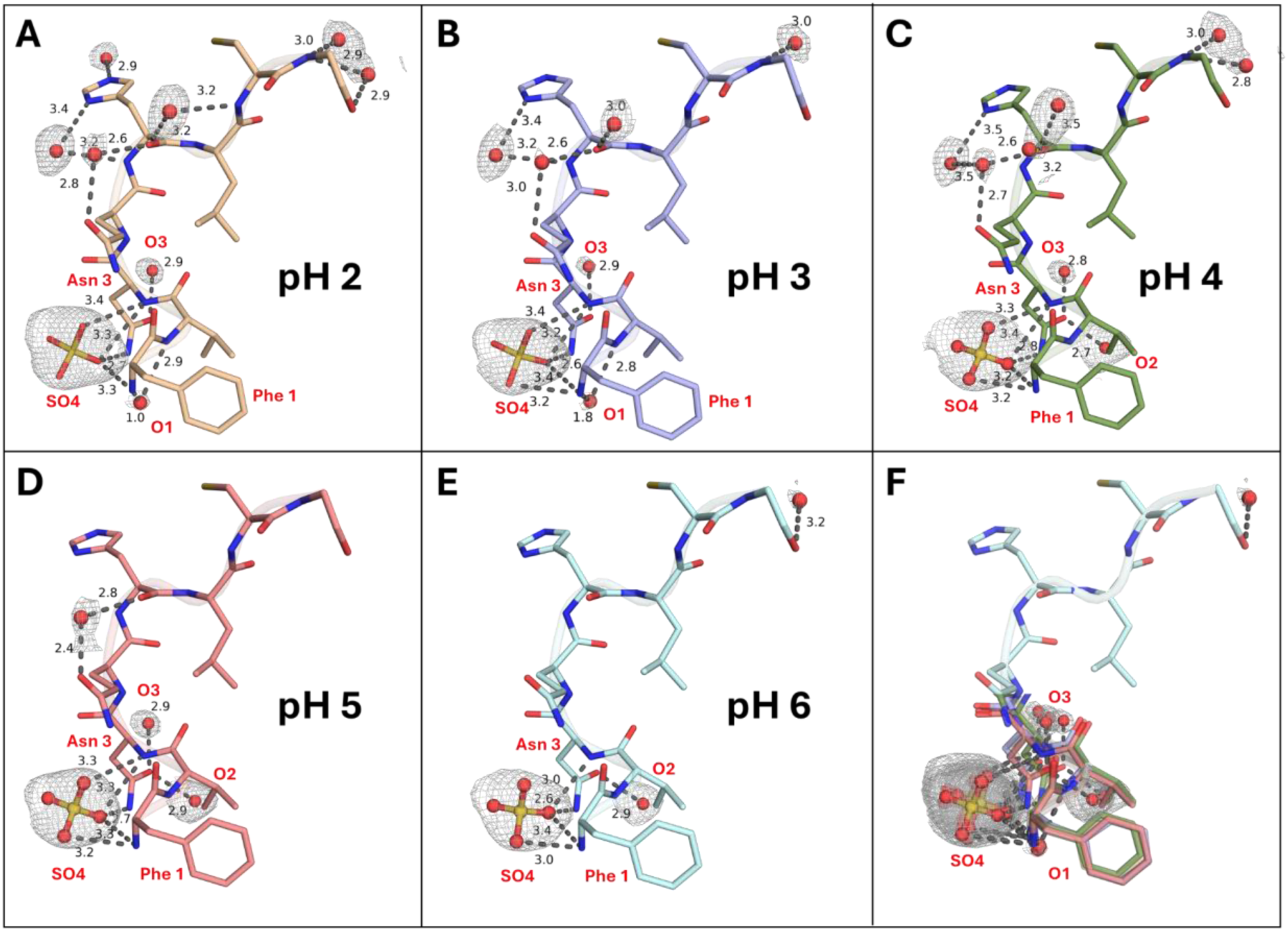
Water and sulfate coordination of key allosteric residues (Phe1 and Asp3) in mIns molecules. **(A, B)** At pH 2 and 3, water molecules O1 and O3, in conjunction with sulfate coordination, stabilize Phe1 and Asp3. **(C, D)** At pH 4 and 5, water molecules O2 and O3, instead of O1, contribute to the stabilization of Phe1 and Asp3, along with sulfate coordination. **(E)** At pH 6, a single water molecule (O2) stabilizes the key allosteric residues in concert with sulfate, indicating a dynamic shift in water coordination as the system approaches the isoelectric point of mIns. **(F)** Superposition of key allosteric residues across the pH range of 2–6 reveals positional shifts in water molecules, revealing pH-dependent rearrangements in the coordination network.

**Supplementary Figure 6.**
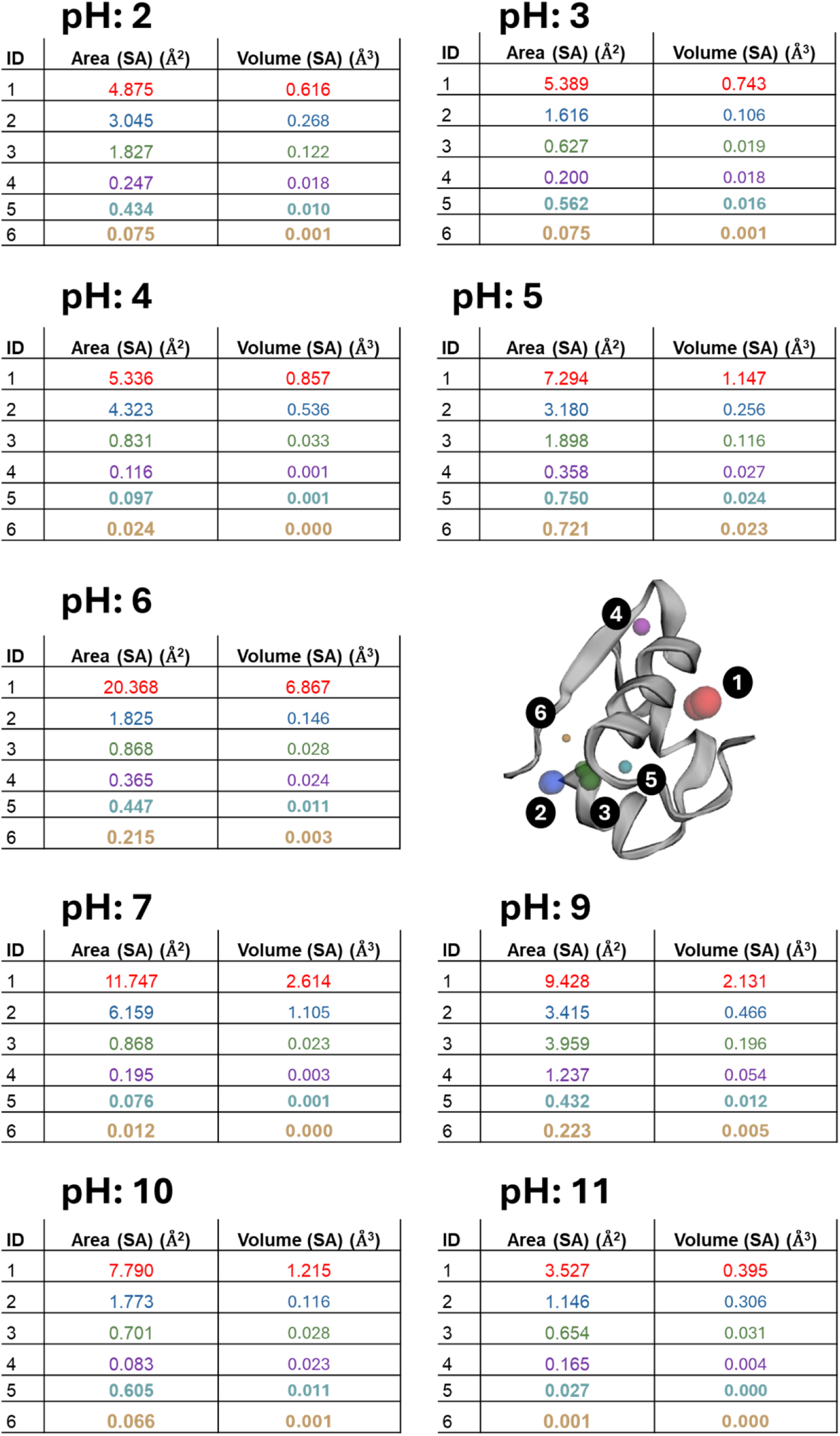
Cavity analysis in the pH range 2–11. The table displays cavity area (Å²) and volume (Å³) measurements at each pH. A total of six cavities were identified in the monomeric structure, ranked by size with cavity ID numbers, and indicated on the monomer structure consistently. The data reveal a pH-dependent cavity remodeling, with the largest cavities observed at pH 6 (Area: 20.368 Å², Volume: 6.867 Å³), corresponding to the isoelectric point where electrostatic repulsion is minimized. In the acidic range (pH 2–5), cavities progressively expand (pH 2: 4.875 Å², 0.616 Å³ → pH 5: 7.294 Å², 1.147 Å³), likely due to increasing charge polarization and solvent interactions. Beyond pH 6, cavities shrink progressively in the alkaline range (pH 7: 11.747 Å², 2.614 Å³ → pH 9: 9.428 Å², 2.131 Å³ → pH 11: 3.527 Å², 0.395 Å³), suggesting that reduction of cavities at extreme pHs is likely attributable to the extensive coordination of structured water molecules and the enthalpically favorable formation of solvent-mediated hydrogen bonding networks ([Djikaev and Ruckenstein, 2013](https://pubs.acs.org/doi/abs/10.1021/jp312631c)) to maintain structural integrity.

**Supplementary Figure 7.**
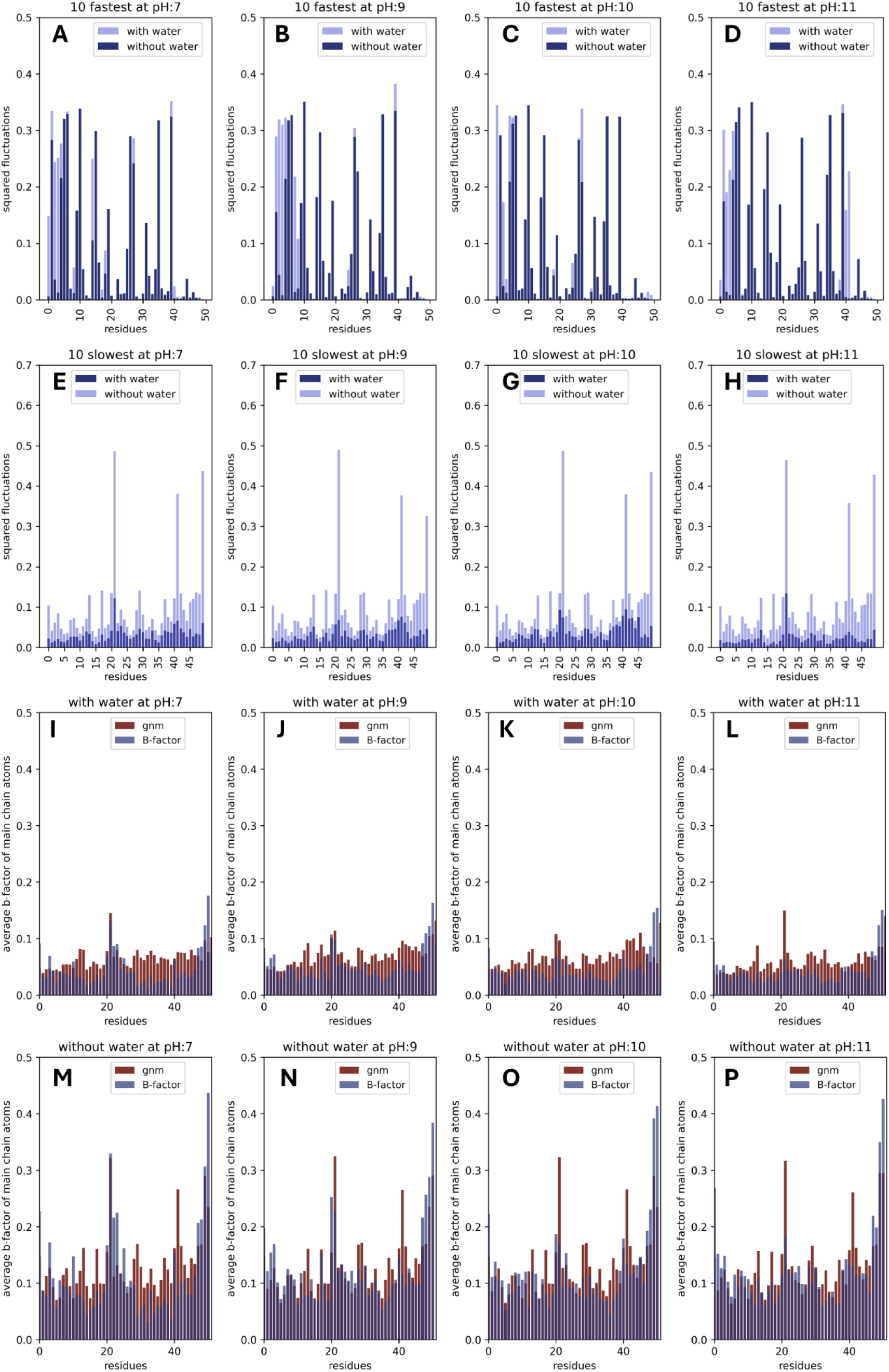
GNM analysis of reference structures (1APH, 2BPH, 3CPH, and 4DPH) at the pH range 7-11. **(A-D)** Local protein motions from the ten fastest GNM modes, comparing structures with and without water coordination in the pH range of 7-11. The mean-square fluctuations of individual modes are plotted for each residue of the structures. **(E-H)** Global protein motions from the ten slowest GNM modes. The weighted squared fluctuations of structure, both with and without water coordination, are presented after normalization. **(I-L)** and **(M-P)** shows the correlation between theoretical GNM fluctuations and experimental B-factors for each structure, with and without water coordination, respectively.

**Supplementary Figure 8.**
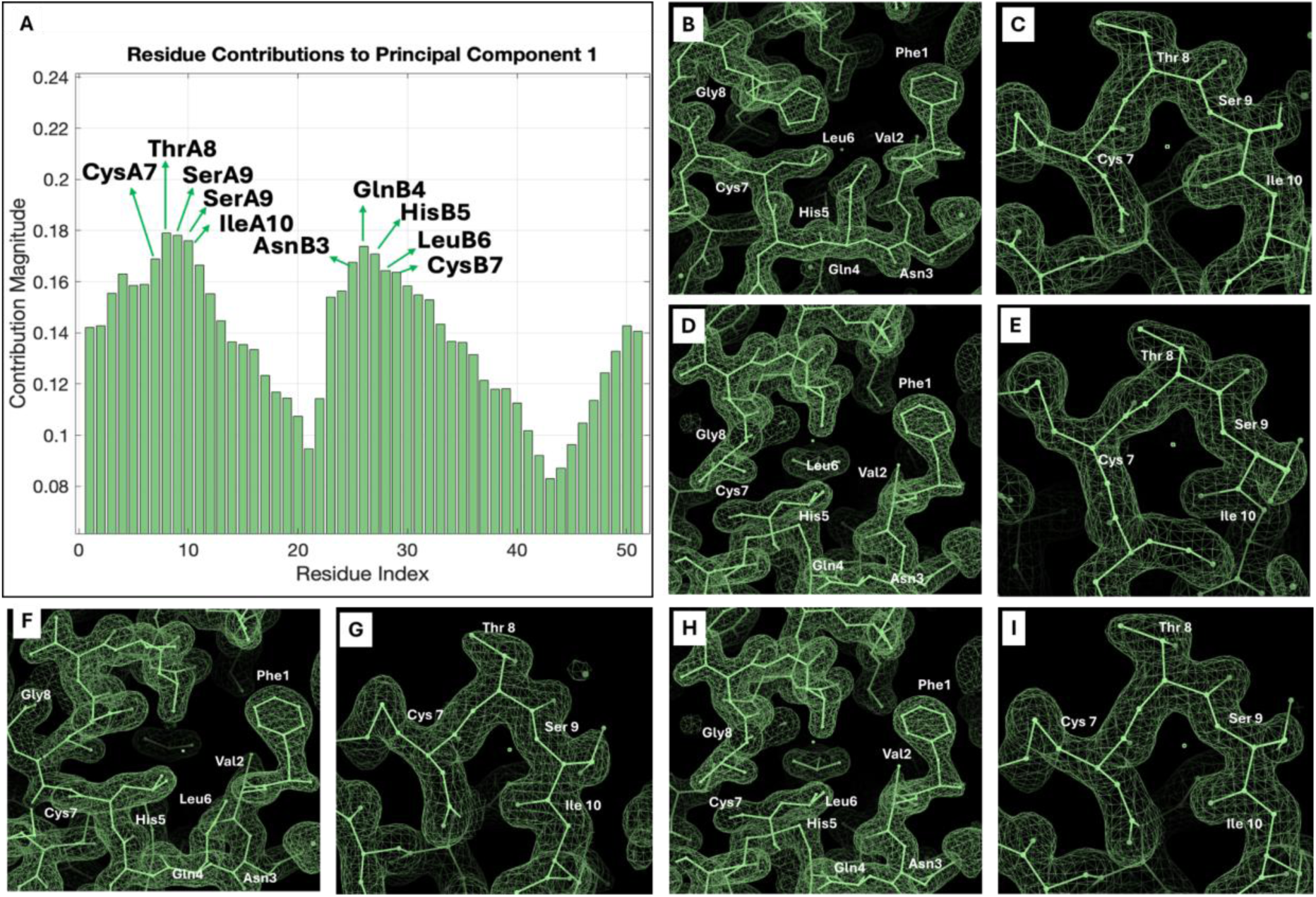
The residue contribution results were derived from PC1, and the electron density of critical residues was also calculated based on PC1. **(A)** Residue contribution analysis identified key residues, including Cys7, Thr8, Ser9, and Ile10 in chain A, as well as Asn3, Gln4, His5, Leu6, and Cys7 in chain B. **(B–I)** The 2Fo-Fc maps contoured at 1σ to reflect full occupancy confirmed the presence of these critical residues for pH 2–6 in each chain.

**Supplementary Figure 9.**
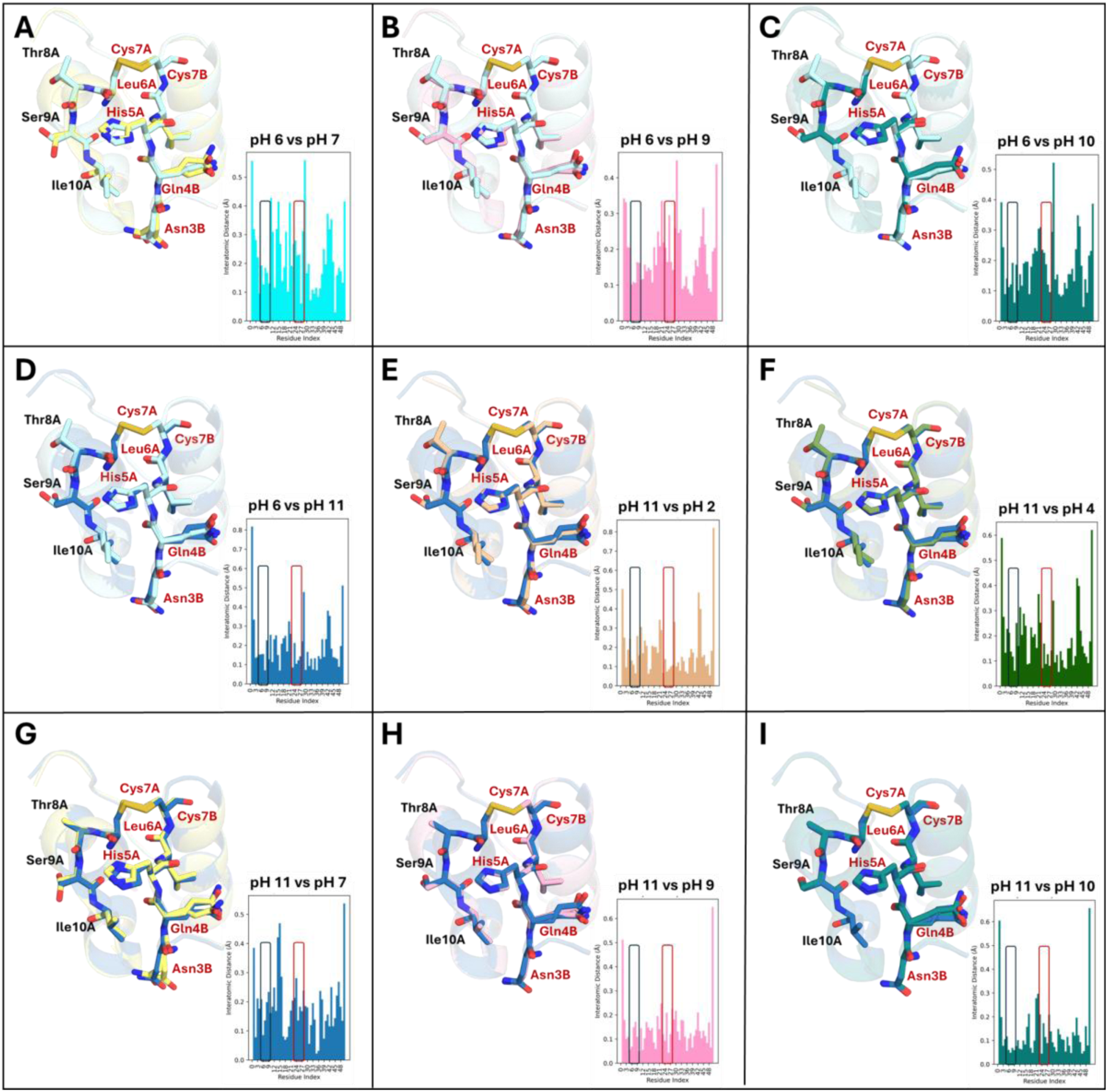
Interatomic distance analysis and structural alignment of outlier structures (pH 6 and pH 11) with cluster 1 (pH 2-5) and cluster 2 (pH 7-10). **(A-D)** Pairwise & structural alignment between pH 6 vs pH 7, 9, 10, and 11 revealed no major conformational changes. **(E-I)** Pairwise & structural alignment between pH 11 vs pH 2, 4, 7, 9, and 10 displayed no major conformational changes. Variant residues between porcine and human insulin are highlighted with a black rectangle, while allosteric sites are indicated with a red rectangle.

